# The spike tip protein of bacteriophage T4

**DOI:** 10.1101/2025.08.28.672839

**Authors:** Yves Mattenberger, Ekaterina S Knyazhanskaya, Mikhail M. Shneider, Sergey A. Buth, Sergey Nazarov, William P. Robins, Petr G. Leiman, Dominique Belin

**Author notes:** Equal contribution (Yves Mattenberger, Ekaterina S Knyazhanskaya).

## Abstract

Contractile injection systems (CISs) – bacteriophage tails, tailocins, and bacterial type VI secretion systems – penetrate the envelope of the target cell by employing a contractile sheath-rigid tube mechanism. The membrane-attacking end of the tube carries a spike-shaped complex that ends with a spike tip. In bacteriophage P2, the spike and spike tip proteins are fused, and we used this phage to show that sheath contraction results in the translocation of the spike into the periplasm of the host cell. In bacteriophage T4, the spike and spike tip proteins are encoded by different genes. We show that the ORFan gene *5.4* codes for the spike tip protein of bacteriophage T4. Using an amber nonsense mutation, we show that the gp5.4 protein is dispensable for bacteriophage T4 particle assembly but essential for bacteriophage fitness and infection of bacteria with truncated lipopolysaccharides.

## Introduction

Contractile injection systems (CISs) are a class of macromolecular machines that utilize a contractile sheath-rigid tube assembly for translocation of proteins and DNA across lipid membranes or simply piercing a hole in the plasma membrane (**Fig. 1A**) [1]. CISs include bacteriophage tails, tailocins (R-type pyocins, *Clostridium difficile* “diffocins”, *Photorhabdus* Virulence Cassette (PVC), *Serratia entomophila* antifeeding prophage (afp), *Pseudoalteromonas luteoviolacea* Metamorphosis Associated Contractile (MAC) arrays and many others), and bacterial Type VI Secretion Systems (T6SS) [2–15]. Using energy stored in protein conformation during assembly, the sheath contracts and drives the tube through the host cell envelope [16]. In phages and tailocins targeting prokaryotes, the tube contains a tape measure protein (TMP) [1], which must exit the tube for it to serve as a channel for DNA and/or ions in the inner membrane. At its membrane-attacking end, the tube carries a central spike-shaped protein or protein complex, which plugs it shut (**Fig. 1A**).

**Figure 1.**
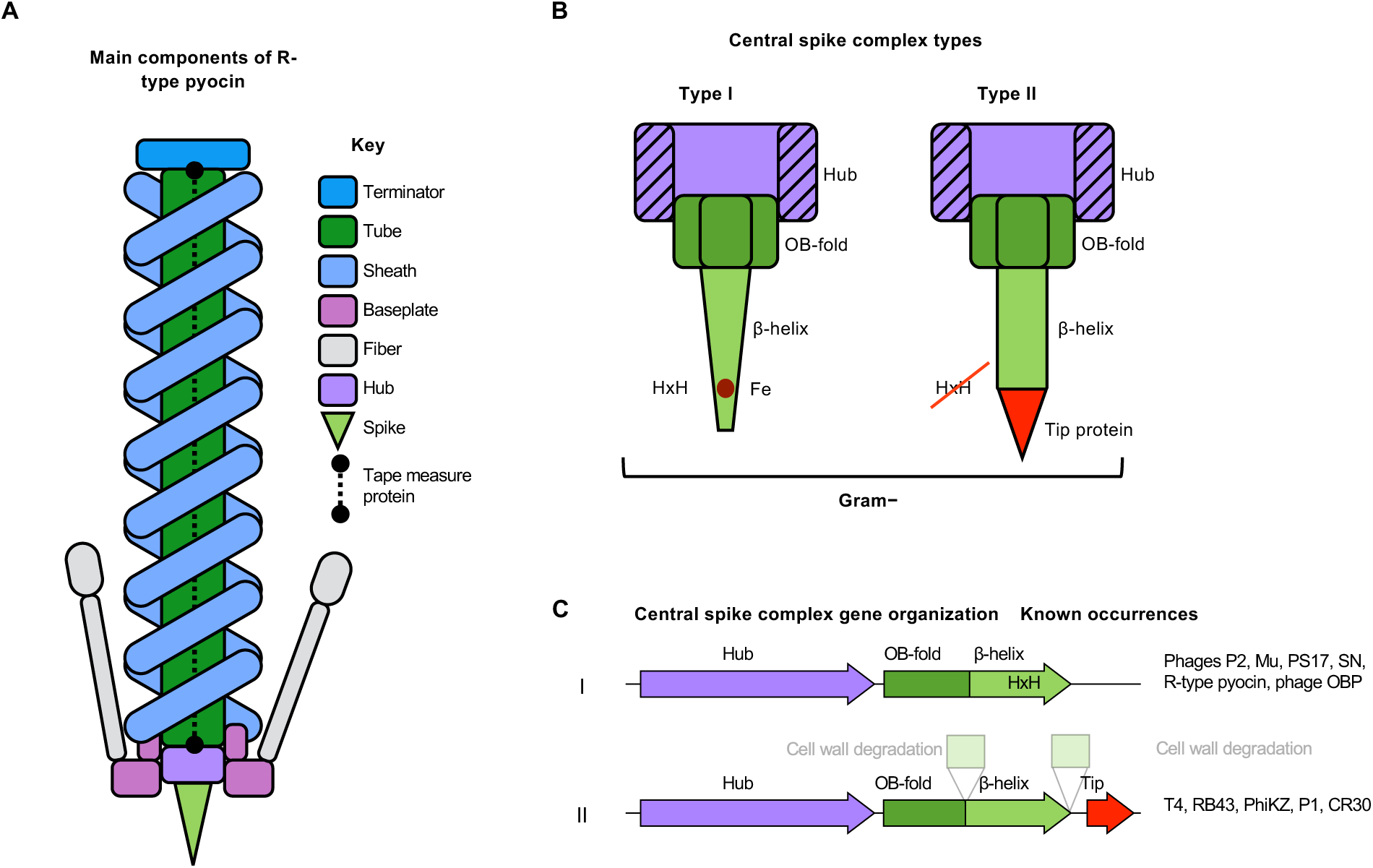
Organization of a contractile injection system and its membrane-attacking spike. **A**, A schematic showing the main components of R-type pyocin, one of the simplest CIS [16]. **B**, Two most common architectures of central spike complexes. For clarity, the diagram does not show facultative enzymatic or partner-attachment domains. The type of targeted bacterial cell wall is indicated. **C**, Gene organization of central spike complexes. The diagram depicts only two of the most common occurrences of facultative domains, which are shown semitransparent. In type II complexes, the tip gene is not necessarily immediately downstream of the spike gene.

Electron microscopy studies of bacteriophages attached to Gram-negative (diderm) bacteria show that upon sheath contraction the tube does not cross the periplasmic space completely. The tube’s tip stops in the periplasm, and the cytoplasmic membrane blebs out to interact with it [17–19]. The fate of the spike complex in this process has been a long-standing question because it determines the site of TMP release and the role of the TMP in DNA translocation.

The spike complex is essential for the assembly of the baseplate and, thus, for the entire CIS machine [20–22]. The spike complex consists of a trimeric hub, a trimeric central spike, and, optionally, a monomeric spike tip (**Fig. 1B**) [1]. The spike contains two domains, an OB-fold and a β-helix. Spike complexes in which spike sequences contain a HxH (histidine, any residue, histidine) motif close to the C-terminus were proposed to be devoid of a tip protein (**Fig. 1C**) [23]. Such spikes are equipped with a pointy apex domain that contains a buried Fe ion coordinated by three copies of the HxH motif in the trimer. The same motif is found at various locations along the length of many host receptor-binding fibers in diverse phages and tailocins (although not near the fiber’s very tip), where it also binds a Fe ion [24, 25]. Spike proteins lacking an HxH motif near their C-terminus have been proposed to carry a tip protein that stabilizes and sharpens the membrane-attacking tip of the spike complex [23]. A majority of known tip proteins belong to the Proline-Alanine-Alanine-aRginine (PAAR) repeat superfamily [26]. These proteins form pointy conical extensions on the T6SS spikes (valine-glycine repeat protein G, VgrG) [21] and can bear effectors covalently attached to it [26].

Spike tip proteins display some of the highest sequence conservation across all CIS proteins. For example, pairwise comparison of proteins comprising the tails of *Escherichia coli* phages T4 and P1 using BLAST [27] results in only one matching pair: T4 gp5.4 matches P1 UpfC with an E-value of 1×10^−8^. Although the high conservation of spike tip proteins suggests that a tip-carrying spike must be evolutionary advantageous, no lethal mutation has ever been reported for a spike tip gene in any bacteriophage.

When the genome of phage T4 was sequenced [28], more than 100 genes encoding proteins of unknown function (ORFans) were identified. The proteins encoded by these ORFans are considered to be non-essential for phage growth under laboratory conditions. All but three expressed from early promoters. The remaining near contiguous ORFans *(5.1, 5.3 & 5.4*) that are expressed from late promoters could be involved in virion assembly or present in viral particles.

This study reports on the structure of the gene product of ORFan *5.4*, a protein localized to the T4 baseplate. A single copy of this protein is attached to the C-terminal end of the gp5 trimer. Here, we defined the physicochemical properties of this attachment. We confirm the identity of the T4 spike tip protein in the phage particle using cryo-electron microscopy (cryo-EM) of a phage carrying an amber mutation in gene *5.4*. We also show that *Escherichia coli* phage RB43, a close relative of T4, carries a unique tip on its T4 gp5-like spike (encoded by *orf204*) that is possibly encoded by gene *orf205w*. To determine the cellular localization of the spike during infection, we used phage P2 for practical reason because its spike and sipke tip are fused in a unique protein, GpV. We localized GpV to the periplasm after infection by bacterial fractionation and western blot, confirming previous suggestion that the cytoplasmic membrane is not pierced during infection.

While T4 gene *5.4* is not essential for phage growth under laboratory conditions, it provides a strong evolutionary advantage. Furthermore, a mutation in the bacterial LPS synthesis gene *hldD* that affects only slightly infections by wild type T4 prevents the growth of mutant phage particles devoid of gp5.4. Our findings reveal the importance of the spike tip proteins for phage infection.

## Results and Discussion

### Crystal structure determination strategies

We used HHpred [29] and synteny of genes encoding structural proteins to identify spike and spike tip proteins with diverse sequences in several phages including T4, RB43, P2, PhiKZ, OBP and SN (**Fig. 1C**). To produce a T4 gp5-gp5.4 complex in soluble form, we used a grafting strategy that worked previously for two T6SS spike tip proteins [26]. We co-expressed gp5.4 with a short fragment of the T4 gp5 spike protein β-helix (residues 484-575). This gp5 fragment, previously called gp5β-BC2 [30], is referred as gp5β here.

The crystal structure of gp5.4 was solved by molecular replacement [31, 32] using gp5β as a search model. The model-free electron density, which extended from the C-terminal end of gp5β, was used to build the atomic structures of tip proteins *de novo*. The interpretation was aided by a 1.15 Å resolution of electron density map of the complex.

### Structure of T4 gp5.4

The fold of T4 gp5.4 is similar to that of T6SS tip proteins c1882 and VCA0105 described earlier [26]. It consists of three β-hairpins wrapped around each other in a right-hand screw-like fashion (**Fig. 2A)**. The hairpins have different lengths, which results in a tapered cone-shaped structure. The length of the hairpin is not related to its position in the amino acid sequence of the PAAR repeat protein: the longest hairpin in T4 gp5.4 is hairpin two.

**Figure 2.**
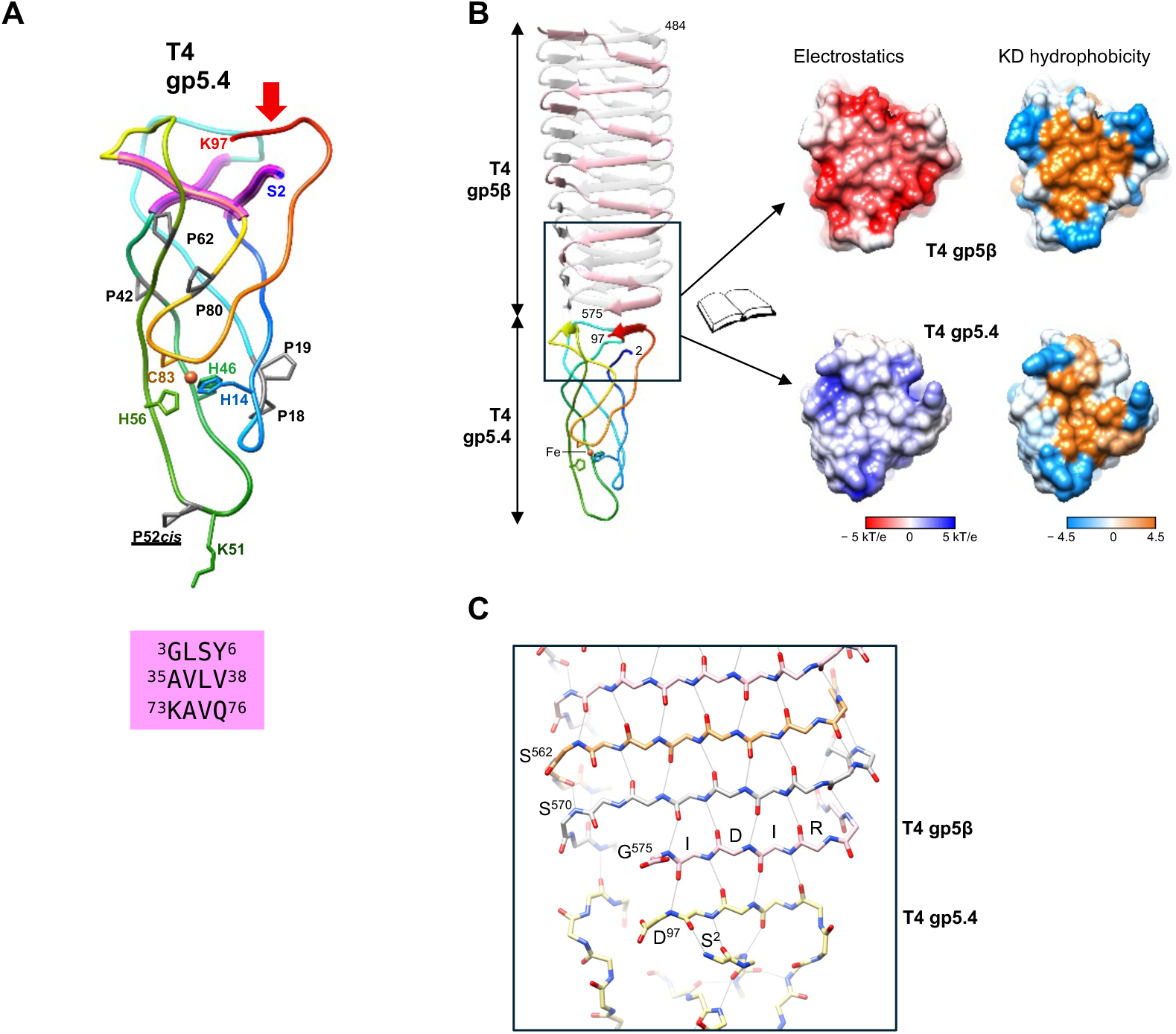
Structure and properties of spike-tip complexes. **A.** Ribbon diagrams of the PAAR repeat tip protein gp5.4 with key residues labeled. The structural elements equivalent to PAAR motif sequences are colored magenta. The corresponding amino acid sequences are given below in the magenta box. The prolines are colored gray. The cyclically permuted element is labeled with a red arrow. **B.** The crystal structure of gp5β-gp5.4 is shown as a ribbon diagram. One of the three chains of gp5β is colored in pink while the other two are in semitransparent gray. The gp5.4 protein is colored in a rainbow pattern along the length of the polypeptide chain with the N-terminus in blue and the C-terminus in red. The molecular surfaces are colored according to their electrostatic potential and Kyle-Doolittle hydrophobicity, respectively. **C.** Hydrogen bond network between main chain gp5β and gp5.4 that seals the hydrophobic interface between two proteins.

The PAAR sequence motif, the defining feature of the eponymous protein superfamilies cd14671, PF05488, DUF4280 or IPR008727, together with two additional copies of identical or similar sequence has been shown to enable a tight, threefold-symmetric crossover of the polypeptide chain in gp5.4 and T6SS spike tip proteins [26]. This property is likely important for giving the monomeric tip protein local threefold symmetry as is required for the interaction with a threefold symmetric spike.

The hydrophobic core of T4 gp5.4 is small and contains only 6-8 residues. It is located between the PAAR repeat crossover region and the metal binding site. The tips of the hairpins are linked by a metal binding site with a tetrahedral coordination sphere, which is formed by three His and single Cys side chains. The identity of the metal was determined as iron (Fe) by X-ray fluorescence spectrometry. The integrity of the metal binding site is important for proper folding because gp5.4 mutant lacking a single histidine (H14A, a His14 to Ala mutation) failed to bind to the gp5 spike.

### The spike-tip interface

T4 gp5.4 is a monomer that binds to the flat tip of the trimeric spike protein β-helix by means of a central hydrophobic patch zipped by hydrogen bonds between the main chains of the spike and the tip protein (**Fig. 2C)** This interface is further enhanced by additional intersubunit hydrogen bonds and ion bridges formed by side chains. This interaction is very similar to that found in the T6SS VgrG-PAAR repeat tip complexes [26] and somewhat similar to the manner by which the monomeric gp38 tail fiber tip protein binds to the trimeric gp37 distal fiber segment in T-even phages [33].

### Location of the spike-tip complex in the tail and confirmation of its identity

Despite the availability of a nearly complete atomic structure of the T4 baseplate [34] [14], the structure of the tip remained unresolved as the tip is located on a symmetry axis while possessing no symmetry itself. To confirm that the cryo-EM density extending from the C-terminal end of the T4 spike gp5 is fully accounted by gp5.4, we created a T4 *5.4* amber mutant T4 *5.4am* and calculated a cryo-EM reconstruction of this mutant grown on the non-permissive *E. coli* strain B178, in which the translation of the *5.4am* gene terminates prematurely at the newly introduced amber stop codon (**Fig. 3A**). The gp5.4 putative density is absent from the cryo-EM map confirming the previous assignment (**Fig. 3C**).

**Figure 3.**
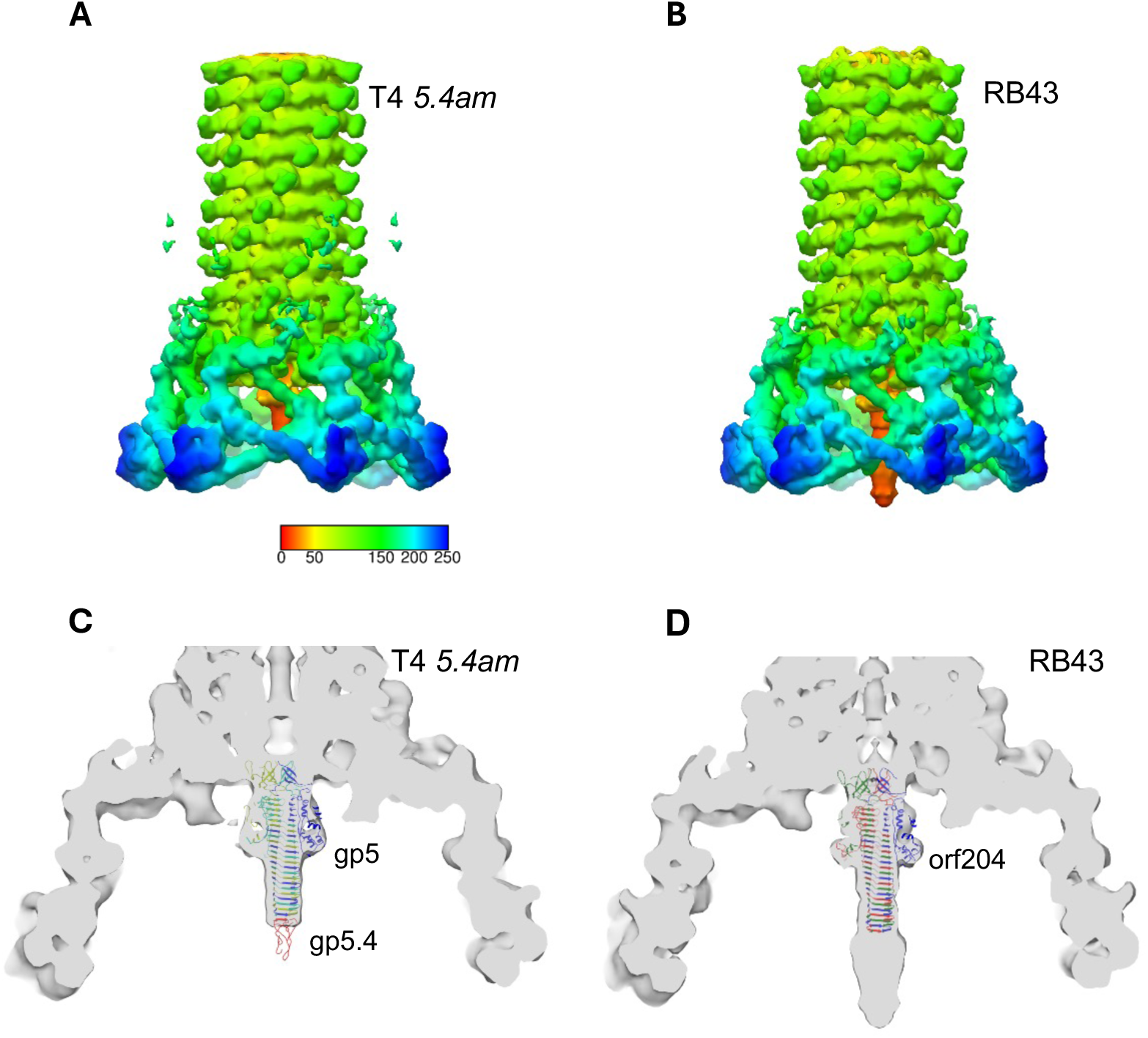
Location of spike-tip complexes in phages. **A** and **B**, Cryo-EM reconstructions of the baseplate regions of T4 *5.4am* (A) and RB43 (B). Isosurfaces colored by cylinder radius with rainbow palette. **C** and **D**, Sliced view of cryo-EM reconstruction of the T4 *5.4am* (C) and RB43 (D) baseplates. The rigid-body fitted (gp27)_3_-(gp5)_3_-gp5.4 protein complex structure, is shown as a ribbon representation fitted into the T4 *5.4am* cryo-EM map (C). The RB43 baseplate with the rigid-body fitted spike (orf204). The structures predicted by AlphaFold are in a ribbon representation.

### The RB43 spike tip protein

Several close relatives of T4 (e.g. phages RB43, RB16, Lw1 [35]) do not contain an ortholog of gene *5.4* in their genome. Although the RB43 and T4 tail proteins show greater than 40% sequence identity, the locus equivalent to T4 genes *5.1* through *5.4* is occupied by a gene (*orf205w*) that encodes a ∼690 residue-long protein in which residues 1-220 are predicted to be an α-helix/β-strand mix, and the rest of the protein to be entirely α -helical. This protein is highly conserved in several phage genomes. Our attempts of producing the RB43 potential spike protein in complex with gp5β were unsuccessful. To better understand the structure of this unusual central spike complex, we calculated a cryo-EM reconstruction of the baseplate region of the RB43 tail. The cryoEM reconstruction of the RB43 baseplate was very similar to that of T4 tail except for the tip region of the spike complex (**Fig. 3B, 3D**). RB43 carries a larger protein at its spike tip. To interpret this map, we used AlphaFold to model the spike (orf204) and the putative spike tip protein (orf205w) of RB43. AlphaFold predicts a model of the RB43 orf204 spike with a very high confidence. The sequence identity of RB43 orf204 to T4 gp5 is 47%, and the two proteins have very similar structures. Similar to T4 gp5, the β-helical domain of RB43 orf204 extends to the very C-terminus of the protein and forms a T4 gp5-like platform at its tip. Although the predicted confidence of the RB43 orf205w spike tip model is low, it is significant, and orf205w can fit in the cryoEM density at the tip of the orf204 spike (Supplementary Fig. 1).

### The spike is translocated into the periplasmic space during infection

To determine the localization of the spike-tip complex in the infected cell, we chose to work with the P2 phage system instead of T4 because the former allowed us to examine the infection process using high multiplicity of infection (MOI), a condition at which T4 is known to cause premature cell lysis known as ‘lysis from without’ or lysis from the outside [36]. Furthermore, as described above, in P2-like spikes the spike and tip are fused into a single gene, gene *V*, so the localization of both functional modules can be studied at once.

We used a gene *V* nonsense mutant of P2 *vir1* (*vir1* stands for lytic) – P2 *vir1* [37, 38] – grown in a permissive host to infect non-permissive *E. coli* C-2 at high MOI, separate the cell lysate into periplasmic, soluble and insoluble cytoplasmic, and membrane fractions, and locate gpV with the help of anti-gpV antibodies (**Fig. 4A**). *E. coli* C-2 does not support growth of P2 amber mutants so no new gpV is made in these cells, and all gpV originates from the infecting phage. GpV was found in the soluble periplasmic fraction (**Fig. 4C**). Even at an MOI of 100, P2 demonstrated a high absorption efficiency, and the titer of unabsorbed phage decreased by two orders of magnitude in 15 minutes post-infection (**Fig. 4B**).

**Figure 4.**
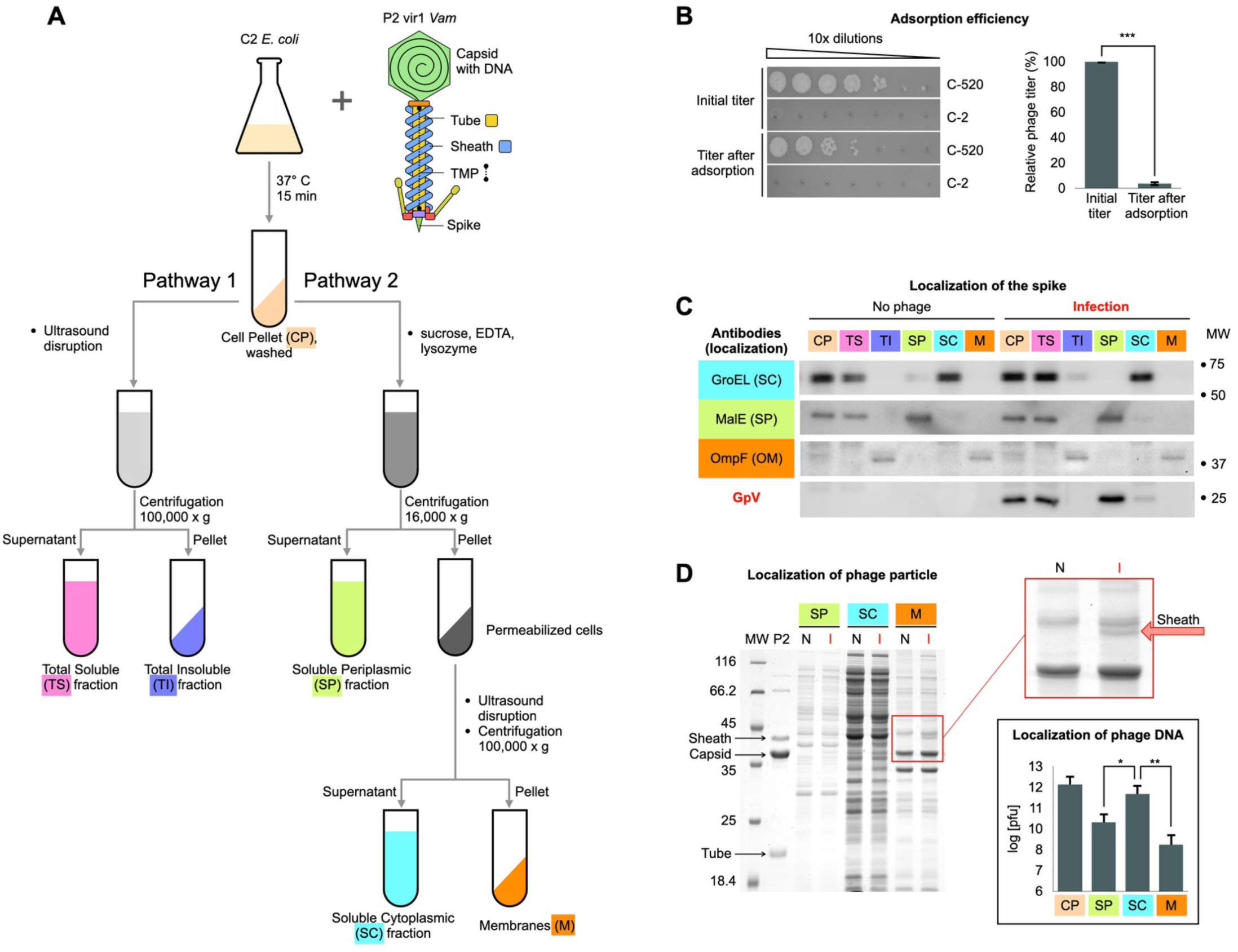
Localization of the P2 phage spike protein in the infected cell. **A,** Flowchart of the fractionation procedure. The same nomenclature (colors and abbreviations) is used to label the cellular fractions in panels C and D. **B**, The two top rows in the left panel show titration of P2 vir1 *Vam46* on either C-520 (permissive, *supD*) or C-2 (non-permissive) *E. coli* lawns. The two bottom rows in the left panel show titration of P2 vir1 *Vam46* that remained in solution after a 15 min incubation with cells. These cells were subsequently used in the fractionation procedure. The right panel shows quantitative representation (mean ± SD) of the titration-presented in the left panel. The experiment was repeated six times with similar outcomes. **C**, Western blot analysis of cell fractions from cultures which were either uninfected (left section of the blot) or infected with P2 Vir1 V*am46* (right section). Each column is a separate fraction of the fractionation procedure shown in panel **A**. The rows correspond to different antibodies used against cellular proteins with known localizations: GroEL is a soluble cytoplasmic protein, MalE (MBP, maltose binding protein) is a soluble periplasmic protein, OmpF is the outer membrane porin F. GpV co-localizes with MalE. The blot is representative of four biological replicates. **D**, A Coomassie stained polyacrylamide SDS gel showing a purified P2 Vir1 phage sample (labeled P2) and fractionated lysates of uninfected C-2 cells (labeled N) or infected with P2 Vir1 *Vam* (labeled I). The red arrow points to the P2 sheath protein; its identity was confirmed by LC/MS/MS analysis. The P2 capsid protein is partially masked by a cellular membrane component with a similar electrophoretic mobility. The inset in a black box (lower right) shows qPCR analysis of P2 genomic DNA (mean ± SD) found in different cellular fractions. The fluorescent signal was converted to the number of plaque forming units (pfus) using a calibration curve as described in the methods section. The experiment was repeated three times with similar outcomes. The significance was determined by Student’s two-tailed *t*-test with one, two, and three stars (*, **, ***) corresponding to p-values of less than than 0.05, 0.001, and 0.0001, respectively.

To extract the periplasmic content, a buffer containing sucrose, EDTA, and lysozyme was used to permeabilize the outer membrane and digest the peptidoglycan (**Fig. 4A**). If this treatment caused the attached phages to dissociate from the cell membrane, a large fraction of gpV would have been wrongly localized. To confirm that the sucrose-EDTA-lysozyme treatment cannot dislodge the phage from the cell membrane, we performed infection at an MOI of 1,000 and attempted to identify the two major components of the phage particle – the capsid and the sheath proteins (gpN and gpFI, respectively) – in the cell pellet. Indeed, both proteins can be identified in the membrane fraction on a Coomassie-stained SDS polyacrylamide gel of the cell lysate (**Fig. 4D**). Because the mobility of the capsid protein was close to that of a major cellular component, we confirmed its identity with mass spectrometry.

Finally, we examined whether the sucrose-EDTA-lysozyme treatment did not interrupt the infection process by causing a premature release of the spike from the tube. This would result in an incorrect localization of the spike in the infected cell. We reasoned that if the spike is released prematurely, the genomic DNA would not reach the cytoplasm of the host cell. We measured the distribution of phage DNA in various cell fractions with the help of qPCR. As shown in the inset in **Fig. 4D**, the amount of phage DNA in the cytoplasm is nearly two orders of magnitude greater than that in the periplasm, and almost four orders of magnitude greater than that in the membrane fraction and similar to that found in the total cell pellet. Thus, the fractionation procedure did not significantly disturb the infection process and did not alter the *in vivo* localization of the spike.

### Gp5.4 increases the fitness of T4

Considering the high degree of conservation and the critical localization of T4 gp5.4-like tip proteins in CISs (**Fig. 1**), we were surprised to find that the absence of functional gp5.4 was not detrimental to T4 in our initial tests. Growth of T4 *5.4am* on non-suppressor (*sup*^+^) *E. coli* appeared to be identical to that of the WT T4+ as measured in a standard growth assay. Neither the plaque morphology nor the intracellular accumulation of virion particles were affected (**Fig. 5A**). The efficiency of plating (EOP) of the mutant phage on *sup*^+^ *E. coli* K12 and B strains was also indistinguishable from that of the WT (Fig. 5D, **Supplementary Fig. 2A**).

**Figure 5.**
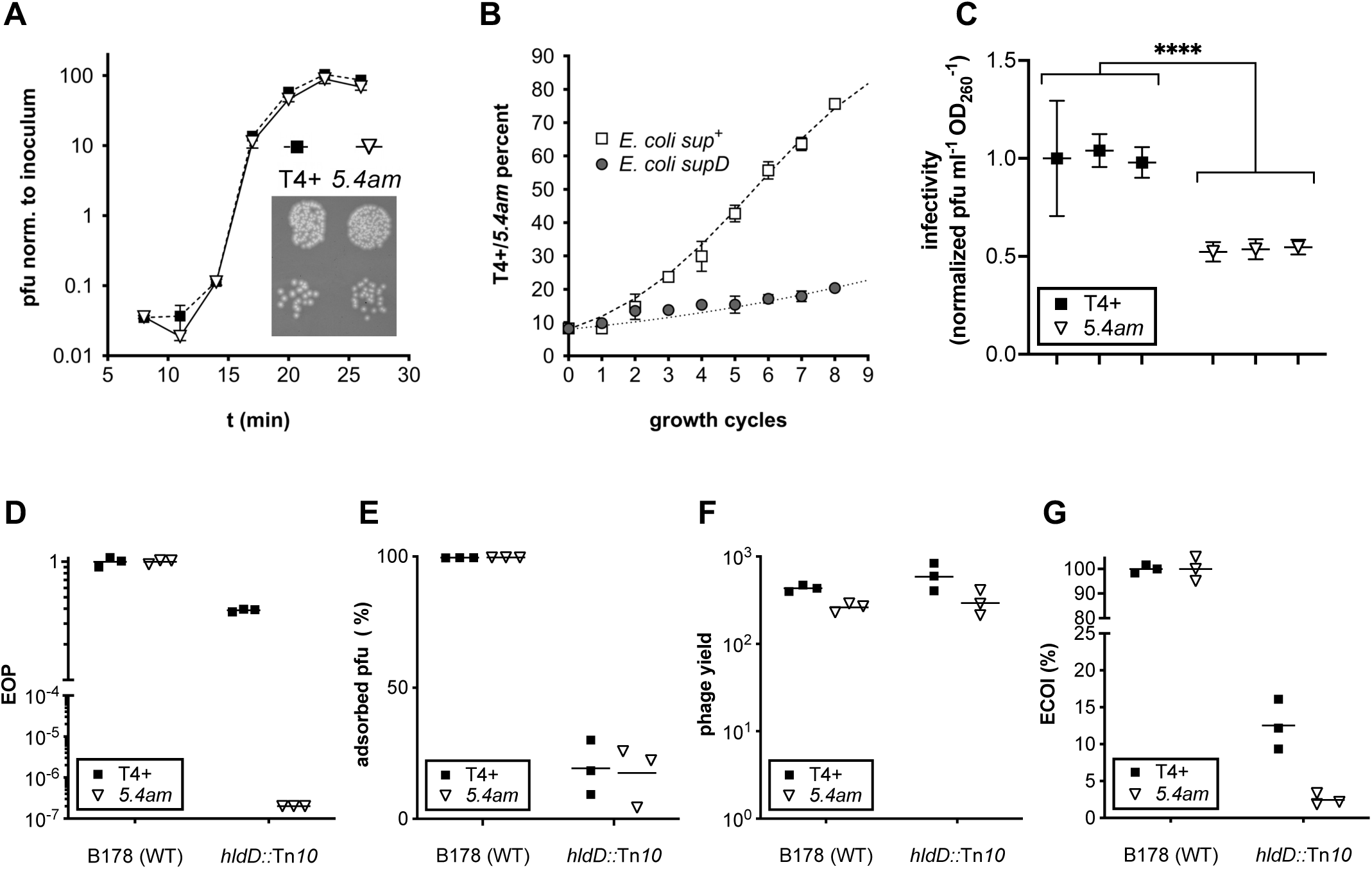
Comparative properties of T4+ and T4 (*5.4am*). **A**, Single growth cycle of T4+ and T4 *5.4am* phages. *E. coli* B^E^ cells (non-permissive = *sup*^+^) were infected at a multiplicity of infection (MOI) of 0.05, and intracellular phage accumulation was measured at the indicated time points. Phage stocks were produced on the *sup*^+^ *E. coli* strain B^E^ and the progenies were analyzed on the *supD* permissive suppressor strain CR63. Data points represent the mean and range of two independent experiments normalized to the input titer. The inset shows the plaque morphology on B^E^ bacteria. **B**, Evolution of the T4+ to T4 *5.4am* ratio when mixed phage inoculums were grown on *E. coli* B^E^ (*sup*^+^) or CR63 (*supD*) cultures over eight successive growth cycles. The initial T4+/T4 *5.4am* ratio was 0.08 and the MOI of each growth cycle was less than 0.1. The percentage of *5.4*^+^ phages was determined by PCR of genomic DNA and restriction. The data points represent the mean and range of two independent experiments. The dashed and dotted lines are mathematical simulations in which the fitness of the T4 *5.4am* mutant phage was 1.55 (*sup*^+^) and 1.14 (*supD*) times lower than that of the WT. **C**, Comparison of the ratio of the infectious to physical virions for T4+ and T4 *5.4am*. The relative ratios of pfu/ml to A_260nm_ are plotted for three independent biological samples of T4+ and T4 *5.4am* which were grown on *E. coli* B^E^ and purified on CsCl gradients (*Materials and Methods*). The titers were determined on *E. coli* B^E^, and the 260nm absorbances were measured by UV spectroscopy. The error bars are SDs which were derived from errors associated with titer and absorbance measurements (each parameter was measured three times). The significance was determined by two-way ANOVA with a p-value of less than 0.0001 (****). **D**, **E, F, G.** Characterization of infection of *sup*^+^ *E. coli* K-12 B178 and its transposon mutant *hldD::*Tn*10* by T4+ and T4 *5.4am*. Efficiency of plating (EOP), adsorption, phage yield, and efficiency of center of infection (ECOI) were determined as described in *Material and Methods*. The cells used for infection were grown to OD_600_ = 0.5 in LB medium supplemented with tryptophan (50µg/ml) and glucose (0.4 %). Phage stocks were produced on non-permissive *sup*^+^ strain B^E^ and purified using CsCl gradients in all experiments except for the phage yield determination studies in which the phage was produced on a permissive *supF* strain (T4-(*5.4am*)-gp5.4 particles). The results represent the mean of three independent experiments.

To determine whether the absence of gp5.4 nevertheless affected the fitness of the phage, we set up competition experiments where the growth of the mutant strain is compared to that of the parental strain over successive growth cycles [39]. We observed that T4 *5.4am* was far less fit than its WT parent T4+ when grown on a non-permissive *sup*^+^ *E. coli* strain. Although the initial phage mix contained 90% of T4 *5.4am* and only 10% of T4+, the latter accounted for more than 70% of the population after eight growth cycles (**Fig. 5B**). A numerical simulation (dashed line in **Fig. 5B**) showed that such results could be obtained if the mutant phage has a 1.55 fitness disadvantage per growth cycle. In a permissive *supD E. coli* strain, which can read through the amber stop codon and efficiently inserts a serine, instead of the tyrosine wild type residue in gp5.4, the growth disadvantage of the T4 *5.4am* mutant was negligible (**Fig. 5B**), confirming that the phenotype was due to the *5.4am* mutation. The small difference could be due to incomplete suppression of the amber mutation by the *supD* suppressor, to several nonsynonymous mutations present in the T4 *5.4am* strain (**Supplementary Table 1**) and/or to the presence of a serine instead of a tyrosine at position 6 of gp5.4. We conclude that the non-essential gene *5.4* nevertheless confers a strong growth advantage to phage T4.

### Gp5.4 influences the infection but not the assembly or stability of the T4 particle

To find the cause of the reduced fitness of T4 *5.4am*, we analyzed the role of gp5.4 in the adsorption to a susceptible host, its influence on the yield of phage progeny during maturation and lysis, and its contribution to the stability of the T4 particle. When stored at 37° C, the infectivity of T4 *5.4am* and T4+ particles as measured by the EOP assay decreased over time in an essentially identical manner (**Supplementary Fig. 2B**) suggesting that absence of gp5.4 does not affect virion stability. The absorption properties of T4+ and T4 *5.4am* measured on non-permissive *sup^+^ E. coli* strains K12 and B^E^ were also similar (**Fig. 5E, Supplementary Fig. 2C**).

To determine the effect of *5.4am* mutation on phage yield independently of its potential effect on host cell binding and DNA injection steps, we compared the burst size of the T4(*5.4am*)-gp5.4 phage – that is a genetically *5.4am* phage equipped with a gp5.4 spike tip, which was produced by growing T4 *5.4am* on the *supF E. coli* strain NM538— with that of T4+. On infection of the non-permissive *sup^+^* strain B178, the burst size of T4(*5.4am*)-gp5.4 was 1.6 times lower than that of T4+ (**Fig. 5F**), showing that the *5.4am* mutation affects either particle assembly or subsequent infection efficiency (adsorption is not affected by absence of gp5.4 cf. above). To resolve this ambiguity, we determined the ratio of infective to physical virion particles for T4 *5.4am* and T4+. The two phages were first produced on the non-permissive *sup*^+^ *E. coli* B178 and purified by isopycnic CsCl gradient centrifugation. The number of infective particles was derived from plaque forming unit (pfu) titers on *E. coli* B^E^, while the number of physical virions was considered to be proportional to the sample’s UV light absorbance at 260 nm as the spectroscopic properties of genomic DNA vastly dominate that of other phage components. For a given absorbance reading, the infectivity of T4 *5.4am* was 2.2 times lower than that of T4+ (**Fig. 5C**). This number is comparable to that of the burst size reduction (**Fig. 5F**), suggesting that the lower burst size is due to a less efficient infection during plating pfu enumeration but not to an assembly defect.

Thus, while gp5.4 does not markedly influence the assembly or stability of the T4 particle, it plays an important role in the infection process without affecting host cell binding properties, suggesting that it acts after phage attachment.

### Deep-rough LPS *E. coli* mutants are resistant to gp5.4-less T4 virions

To further decipher the function of gp5.4, we used transposon mutagenesis to isolate three mutants of the non-permissive *sup*^+^ *E. coli* B178 that were resistant to T4 *5.4am* and sensitive to T4+, thus possessing a gp5.4^−R^-T4^S^ phenotype (**Fig. 6A**). In all three gp5.4^−R^-T4^S^ mutants, the transposon mapped to the same 114^th^ codon of gene *hldD* in the *hldD*-*rfaFC*-*waaL* operon, which encodes four enzymes that drive the biosynthesis of the inner core of the lipopolysaccharide (LPS) (**Fig. 6B**). The product of the *hldD* gene catalyzes the last step of the synthesis of the ADP-L-glycero-β-D-manno-heptose that is then covalently linked to the KDO-Lipid A region by ADP-heptose:LPS heptosyltransferase I (HepI), which is encoded by *rfaC* gene [40] [41], [42]. All three *hldD*::Tn10 mutants exhibited a deep-rough LPS profile and phenotype (**Fig. 6C, Supplementary Fig. 2D**), as expected for mutants lacking the heptoses of the inner core [43].

**Figure 6.**
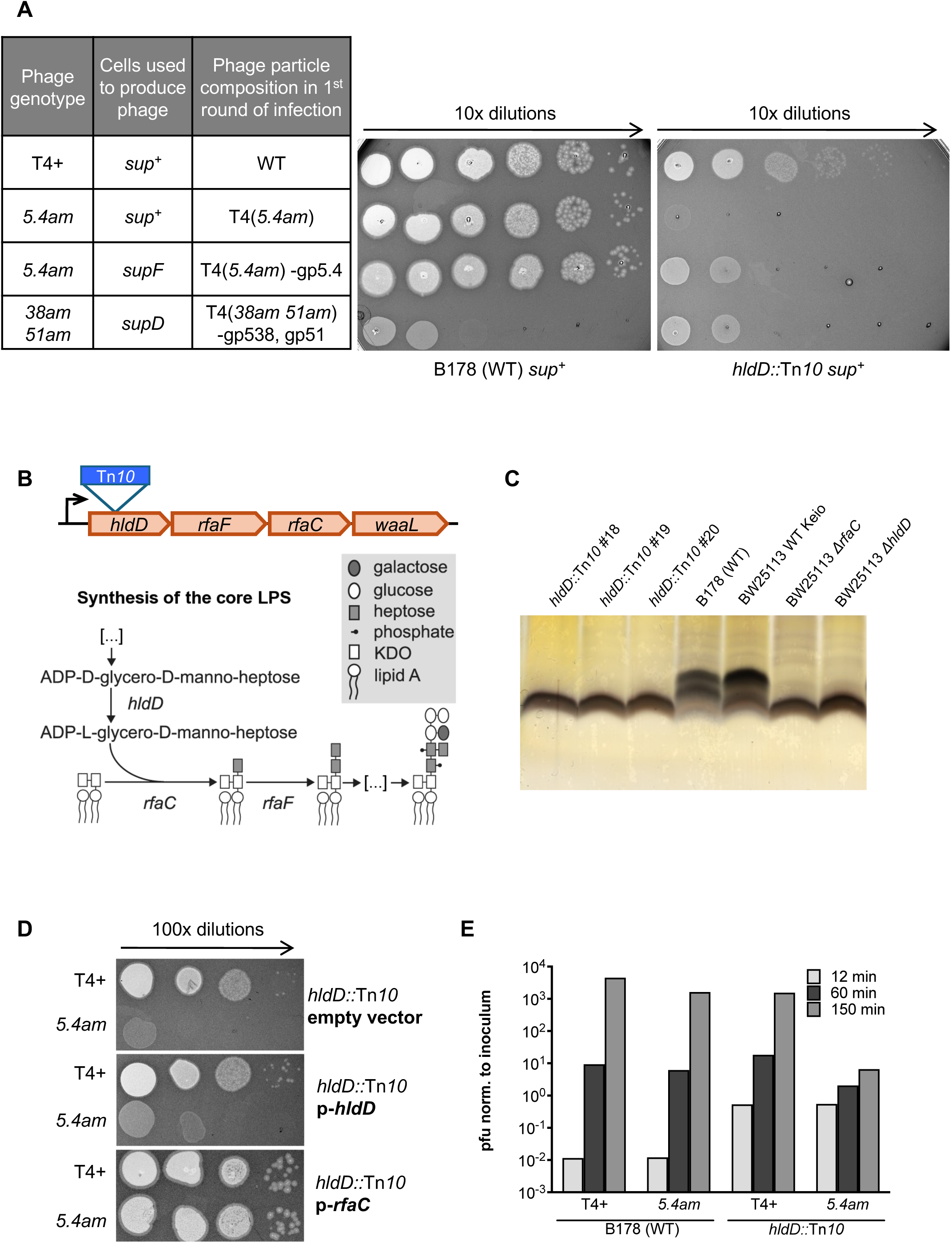
Gp5.4 is essential for T4 growth on deep-rough LPS mutants. **A**, Growth of T4+, T4 *5.4am,* and T4 K10 *38am 51am* on B178 (WT) and *hldD::*Tn*10* bacteria. Phage stocks produced on *sup*^+^ (B^E^), *supF* (NM538) or *supD* (CR63) bacterial strains were diluted and spotted onto the indicated bacterial lawns. Results are representative of three independent experiments. **B**, The *hldD*-*waaL* operon of *E. coli* K12 and the biosynthesis pathway of the core LPS, adapted from [61]. **C**, Silver-stained Tris-tricine SDS-PAGE analysis of LPS extracted from the indicated strains. **D**, Growth of T4+ and T4 *5.4am* on *hldD::*Tn*10* transformed with an empty vector (pCA24N) or plasmids (p-) expressing *hldD* (pCA24N-*hldD*) and *rfaC* (pCA24N-*rfaC*). Top agar plates were inoculated with exponentially growing cultures induced with 100µM IPTG for 1 hour and spotted with dilutions of phage stocks produced on a *sup*^+^ *E. coli* strain. Results are representative of four independent experiments. **E**, Adsorption and growth of T4+ and *T4 5.4am* on B178 (WT) and *hldD::*Tn*10* b bacteria infected at a low MOI (between 0.025 and 0.05) at 37°C (see *Materials and Methods*). Aliquots were withdrawn at indicated times, treated with chloroform, and plated on *E. coli* B^E^ at appropriate dilutions.

To understand how the transposon insertion is responsible for the gp5.4^−R^-T4^S^ phenotype, we conducted a series of additional experiments. First, spot assays of the WT T4+ and T4 *5.4am* phages were performed on *E. coli* strains harboring in frame deletions of each gene of the *hldD*-*rfaFC*-*waaL* operon (**Supplementary Fig. 2E**). None of the mutants reproduced the phenotype of the *hldD*::Tn10 mutant exactly. No difference between the EOP of the WT T4+ and T4(*5.4am*) grown on the *rfaF* and *waaL* mutants was detected. The deletion of *hldD* caused only a small reduction of the T4 *5.4am* mutant’s EOP but did not affect the WT T4+. On the other hand, the deletion of *rfaC* made the cells completely resistant to T4 *5.4am* and significantly reduced the EOP of the WT T4+. Furthermore, plasmid expression of *rfaC* but not *hldD* reversed the gp5.4^−R^-T4^S^ phenotype of the *hldD*::Tn10 mutant (**Fig. 6D**). We concluded that the transposition event in *hldD* exerts a polar effect on *rfaC* which severely reduced the expression of RfaC/HepI.

Thus, the gp5.4^−R^-T4^S^ phenotype was due to a complete inactivation of HldD and a strong reduction of RfaC expression. It has been demonstrated that the LPS molecules produced by *rfaC* and *hldD* null-mutants differ by a single heptose residue: an *rfaC* null mutation results in a heptoseless LPS that terminates with a KDO residue, while LPS molecules of the *hldD* null-mutant contain one additional atypical heptose isomer ADP-D-glycero-ß-D-manno-heptose [44, 45]. The phenotype difference between the *rfaC* deletion mutant (**Supplementary Fig. 2E**) and the polar *hldD*::Tn10 mutant (**Fig. 6D**) was probably due to a minor fraction of heptose-containing KDO_2_-lipidA in its outer membrane.

### Gp5.4 is required for DNA injection during infection of deep-rough LPS mutants

To understand the reason for the specific resistance of *hldD*::Tn10 bacteria to T4 *5.4am*, we characterized the infection process of *hldD*::Tn10 mutants by T4 *5.4am* and T4+ using the same set of experiments as for the WT *E. coli* B178 (**Fig. 5D to 5G**).

The EOP of T4+ on the *hldD*::Tn10 *E. coli* mutants was slightly reduced (**Fig. 5D and 6A**). The EOP of T4 *5.4am* on the same mutants was strikingly low, at the level of the *5.4am* mutation reversion rate (less than 2 x 10^-7^). Similar results were obtained in liquid culture: after several uncoordinated growth cycles (150 min) on *hldD*::Tn10, T4 *5.4am* yielded only 6 pfu per initial infecting phage, 240-fold less than T4+ (**Fig. 6E**). The adsorption efficiency was reduced by about 80% for both the WT T4+ and T4 *5.4am* (**Fig. 5E**). T4 has been shown to be able to adsorb to *E. coli* in both an OmpC-dependent mode and an OmpC-independent mode [46] [47]. In the former mode, T4 adsorption is unaffected by the LPS outer core composition but is strongly reduced in mutant with LPS shorten to the inner core (deep-rough mutations), a phenotype that could be explained by the greater than 90% reduction in porin levels, including OmpC, in deep-rough mutants [48]. In the OmpC-independent mode, T4 adsorption is restricted to LPS whose outer core are capped by a terminal, unbranched glucose [46] [47], as exemplified by the complete absence of growth of the WT T4+ on a *ΔompC ΔrfaC* mutant (**Supplementary Fig. 2E**). While absence of gp5.4 did not change T4 adsorption efficiency on the *hldD*::Tn10 mutant, our results confirm that deep-rough LPS mutation strongly reduced T4 binding to *E. coli*. Finally, the burst size of T4(*5.4am*)-gp5.4 on the *hldD*::Tn10 *E. coli* mutant was about 2 times smaller than that of the WT T4+ (**Fig. 5F**), which is consistent with the reduced infectivity of T4 *5.4am* (**Fig. 5C**).

To test whether the *5.4am* mutation affects DNA delivery during infection, we used the efficiency of center of infection assay (ECOI, also called transmission coefficient). Typically, the ECOI assay evaluates the production of at least one viable phage progeny after plating the infected cells on a permissive host. As our previous experiments show that the *5.4am* mutation does not affect particle assembly or host cell binding properties, ECOI should probe the number of successful DNA delivery events. We found that the ECOIs of T4+ and T4 *5.4am* on *E. coli hldD*::Tn10 were 8 and 20 times lower, respectively, than those on the WT *E. coli* B178 (**Fig. 5G**). While the T4+ ECOI is indicative of a less efficient adsorption of the phage particle to the LPS-mutant host cell surface (both reversible and irreversible attachment), the much-reduced ECOI of T4 *5.4am* compared to T4+ shows that the DNA delivery of the T4 *5.4am* genome is severely impaired.

This conclusion is further supported by comparing the infection of *hldD*::Tn*10* bacteria by T4 *5.4am* viral particles equipped with functional gp5.4 (T4(*5.4am*)-gp5.4 particles produced on a *supF* strain), by T4 *5.4am* viral particles without gp5.4 (produced on a *sup^+^*strain), and by T4 K10(*38am 51am*) [49] viral particles (produced on a *supD* strain) (**Fig. 6A**). As expected T4 *5.4am* particles without gp5.4 could not form plaques on the LPS mutant bacteria while they produced normal plaques on a wild-type bacterial host. When equipped with gp5.4, the T4(*5.4am*)-gp5.4 viral particles still could not form plaques on the LPS mutant, although they did significantly kill the bacteria at high concentrations. This is similar to the effect observed with high concentrations of T4 K10(*38am 51am*) viral particles, which cannot propagate on *sup*^+^ strains because amber stop mutations in tail (*51am*) and long tail fiber (*38am*) genes prevent the assembly of infectious particles. This suggests that T4(*5.4am*)-gp5.4 particles were able to infect and kill LPS mutant bacteria but not to produce a progeny that could further infect these bacteria. Taking into account the facts that gp5.4 is not essential for T4 viral particle assembly (**Fig. 5F**), that the *hldD*::Tn*10* mutant supports the production of T4+ and T4 *5.4am* particles (Fig 5F), and that adsorption of T4 virion on *hldD*::Tn*10* bacteria is not strongly affected by gp5.4 absence (**Supplementary Fig. 2C**), it thus follows that the absence of gp5.4 after the initial infection must have severely impaired the DNA delivery step from the T4 *5.4am* progeny particles into *hldD*::Tn*10* bacteria in the following infections.

Combining the data from all the experiments described above with the critical position of gp5.4 at the tip of the membrane-attacking spike, we conclude that gp5.4 facilitates the penetration of the cell envelope by the phage tail. The results of our experiments also suggest that the interaction between phage fibers and shortened LPS of *hldD*::Tn10 might be insufficient to hold the phage particle with a blunt gp5.4-less spike in place for membrane piercing.

### In trans complementation of T4 5.4am with recombinant gp5.4 restores the WT phenotype

As a final test we asked whether the gp5.4 function could be provided in *trans*, i.e. whether the defective phenotype of T4 *5.4am* on the restrictive *hldD::*Tn*10* strain could be restored by the expression of gp5.4 from a plasmid in the bacterial strain used to produce the phage.

*T4 5.4am* requires a functional gp5.4 to grow on *hldD::*Tn*10* (**Fig. 6A**). The function is restored in *supF* cells. We transformed the *hldD::*Tn*10* strain with a plasmid expressing the *supF* allele under the control of a late T4 promoter (p-*supF*). We then used T4+ and T4 *5.4am* grown in a *supF* strain –thus carrying the WT spike tip (a T4(*5.4am*)-gp5.4 particle)–to infect the bacterial *hldD::*Tn*10 supF* mutant to verify that the two phages have identical phenotypes (**Fig. 7A, columns 1 and 2**). As a nonpermissive control, we infected *hldD::*Tn*10* harboring pBR322 ΔTc (empty vector) and observed the original restrictive phenotype (**Fig. 7B columns 1 and 2**). Thus, in the *hldD::*Tn*10 supF* background, the EOP fully depended on the composition of the particles that were used for the initial infection event, independently of their genotype, i.e. gp5.4-equiped particles can infect and form plaques on this strain even if they are genotypically *5.4am* while gp5.4-less particles are not able to grow on it.

**Figure 7.**
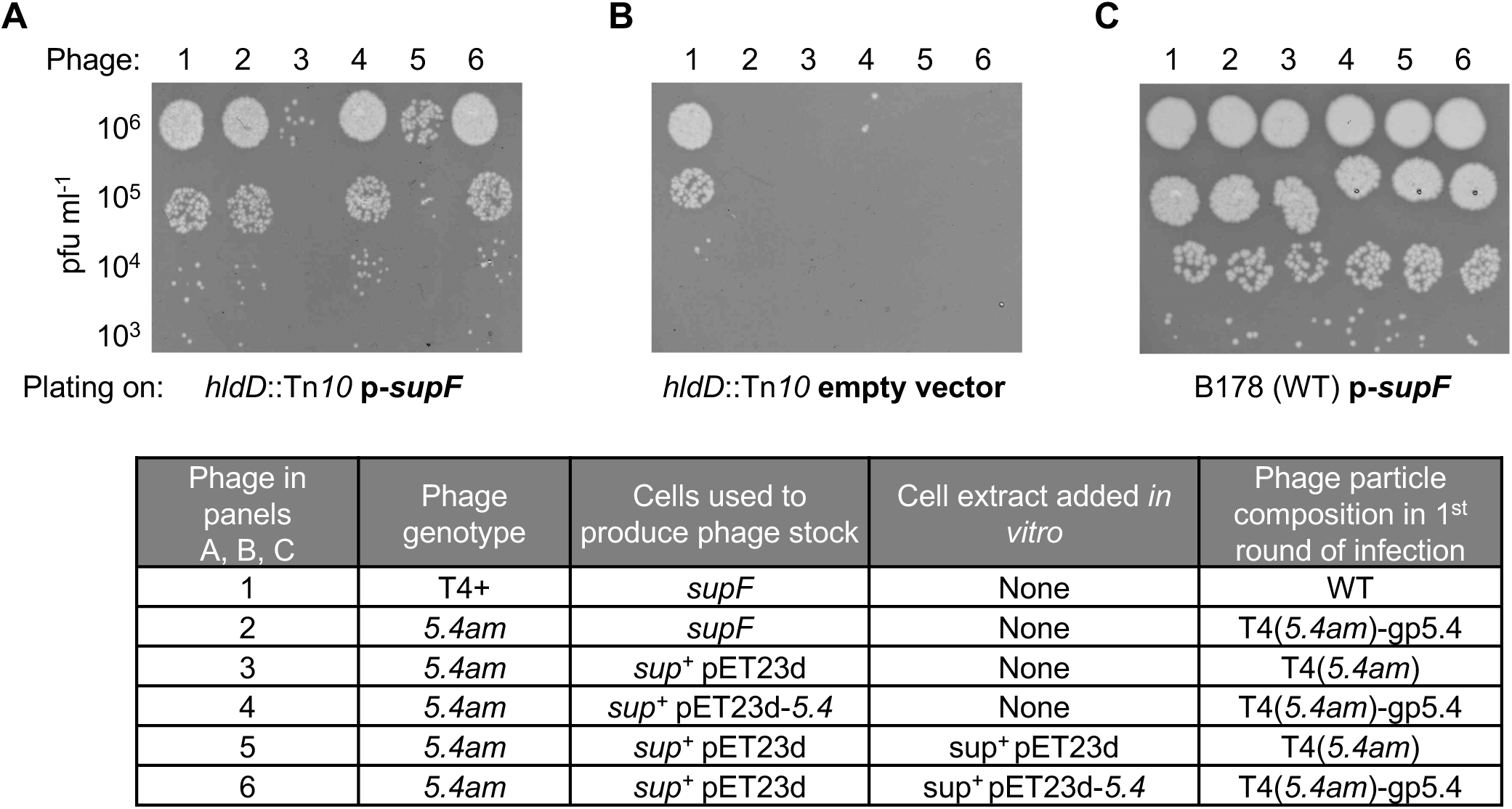
The *5.4am* null mutation can be complemented by providing gp5.4 either *in vivo* during phage growth or *in vitro* to a phage stock. T4+ and *T4 5.4am* phage stocks were either produced on a *supF* strain (*E. coli* K-12, NM538) or on *sup*^+^ strain (*E. coli* B strain, Rosetta DE3) harboring either an empty vector (pET23d) or a plasmid expressing *5.4* (pET23d-*5.4*) induced with 1 mM IPTG. Phage stocks 5 and 6 were complemented with recombinant gp5.4 *in vitro* by incubation for 45 min at 37°C with a fresh cell extract made from uninfected bacteria expressing gp5.4 (see *Materials and Methods* for details). Phage stocks were titrated on *E. coli* B^E^ and equal amounts were spotted onto the *hldD*::Tn*10* harboring pBSPL0+, a plasmid expressing a *supF* suppressor tRNA under the control of a T4 late promoter (p-*supF)*. The phage stocks were also spotted on the *hldD*::Tn*10* mutant harboring pBR322 ΔTc (empty vector) and on B178 (WT) harboring pBSPL0+ (p*-supF*). The experiment is representative of two independent experiments.

We then grew T4 *5.4am* in Rosetta (DE3) cells expressing gp5.4 from an IPTG-inducible pET23d vector to test whether T4(*5.4am*)-gp5.4 particles were produced. As a control, we grew *T4 5.4am* in the same cells harboring an empty pET23d vector. The phages produced on these two strains were then plated on *hldD::*Tn*10* p-*supF* cells. Phages produced on bacteria expressing gp5.4 from the pET23a-*5.4* plasmid had a WT-like phenotype while phage produced on bacteria with the empty vector did not grow (**Fig. 7A, columns 4 and 3, respectively**). Plating these phages on *hldD::*Tn*10* showed that the frequency recombination between the gp5.4 plasmid and phage genome in Rosetta (DE3) cells is about 10^−3^ (**Fig. 7B column 4**), which means that 99.9% of phage population maintained the *5.4am* mutation and the complementation took place at the protein level. We confirmed this finding by sequencing the *5.4* region. We conclude that plasmid-produced gp5.4 could be incorporated in the virions forming T4(*5.4am*)-gp5.4 particles.

Finally, we examined whether gp5.4 can be incorporated *in vitro* into an otherwise fully assembled T4 phage particle. T4 *5.4am* was first produced on a *sup*^+^ bacteria and then incubated with a lysate of Rosetta (DE3) cells containing recombinantly expressed gp5.4. The resulting particles had a WT-like phenotype on *hldD::*Tn*10 supF* bacteria (**Fig. 7A, column 6**), indicating successful incorporation of gp5.4 into the virions. This finding further supports experiments described above showing that gp5.4 does not play a critical role in tail or viral particle assembly despite the strong evolutionary fitness it confers to bacteriophage T4.

## Materials and Methods

### Phage and E. coli strains, plasmids and growth conditions

The phage strains, bacterial strains and plasmids used in this study are listed in **Supplementary Table 2**. The construction of the phage and bacterial strains are described in supplementary Materials and Methods. Unless otherwise stated, all *E. coli* strains were cultivated at 37 °C in LB medium [50] supplemented, for phage experiments, with tryptophan (50 µg ml^-1^) and glucose (0.4 %). For growth on solid medium, 1.5% bacteriological agar was included. Hershey agar (HA, bottom or top agar) plates were used to isolate and count phage plaques [51]. Antibiotics were used at the following concentrations: ampicillin (Ap), 200 μg ml^-1^; chloramphenicol (Cm), 30 μg ml^-1^; kanamycin (Kn), 40 μg ml^-1^; and tetracycline (Tc), 7.5 μg ml^-1^.

### Identification of spike and spike tip proteins with the help of bioinformatics

HHpred [29] and synteny of genes encoding structural proteins was used to identify spike and spike tip proteins in phages T4. The spike protein of phage P2 [23] (**Fig. 1**) contains an HxH (histidine-any residue-histidine) motif near its C-terminus. The spike protein of phage RB43 (gp5) displays 48% overall sequence identity to T4 gp5 spike although the similarity drops off sharply towards the C-terminus and the last 60 residues of RB43 and T4 gp5 have only 15% sequence identity. However, the overall structure of RB43 gp5 is likely to be very similar to that of T4 gp5 with the β-helix extending to the very C-terminus (although the RB43 β-helix is one strand longer). The putative tip protein of RB43 is encoded by gene *orf205w* and it is different to any other tip identified so far.

### Design of construct for expression of gp5-gp5.4 complex

To obtain the structure of gp5.4 in its native context (bound to the gp5 spike), we chose to work with the smallest well-behaving fragment of gp5 β-helix, which we identified in our previous work and called gp5β-BC2 [30]. For brevity, this fragment is named gp5β here. A complex of gp5β-gp5.4 was then purified and crystallized. Gp5.4 did not stain by Coomassie, but the gp5β-gp5.4 complex had a unique elution profile on the MonoQ ion exchange column, and gp5.4 could be detected in the sample with the help of mass-spectrometry.

### Gp5β-gp5.4 complex expression and purification

The pEEva2 plasmid carrying gp5β-gp5.4 DNA complex was transformed into the BL21 (DE3) strain of *E. coli*. The transformed cells were grown at 37 °C in the LB medium, supplemented with ampicillin at the concentration of 200 μg/ml until the optical density reached 0.6-0.8 OD at 600 nm. The medium was cooled on ice to 18-20 °C followed by induction of the gp5β-gp5.4 complex expression by addition of IPTG to a final concentration of 1 mM. After approximately 16 hours at 18 °C overnight, the cells were harvested by centrifugation at 5180 *x g* at 4 °C. The cell pellet was resuspended in 1/50^th^ of the original cell volume in a 20 mM Tris-HCl pH 8.0 buffer supplemented with 300 mM NaCl, 5mM imidazole and 0.02% NaN_3_. The cells were lysed by sonication. The cell debris was removed by centrifugation at 35000 *x g* for 15 minutes at 4 °C. The supernatant was loaded onto the Ni^2+^-precharged 5ml GE HisTrap FF Crude column (GE Healthcare Life Sciences), which was equilibrated with 20 mM Tris-HCl pH 8.0, 300 mM NaCl buffer A. Protein was eluted with 20 mM Tris-HCl pH 8.0, 300 mM NaCl, 250mM imidazole buffer B using two-step gradients employing an AKTApurifier 100 system (GE Healthcare Life Sciences): 1) 15% buffer B (37.5 mM imidazole) – to remove nonspecific bounded proteins and 2) 100% buffer B – actual protein complex elution. The fractions of the elution peak containing the gp5β-gp5.4 complex were pooled and dialyzed overnight with simultaneous TEV His-tag cleavage against 10 mM Tris-HCl pH 8.0, supplemented with 1ml of TEV-protease at the concentration of 1mg/ml, 3 mM DTT, 1.5 mM EDTA for His-tag removal. Digested protein was further purified with ion-exchange chromatography using GE Mono Q 10/100 GL column (GE Healthcare Life Sciences) connected to an AKTApurifier 100 system in 20 mM Tris-HCl pH 8.0 buffer using 0 to 1 M NaCl linear gradient. Selected fractions of the ion-exchange chromatography were analyzed on SDS-PAGE gel. The fractions containing the gp5β-gp5.4 complex were pooled and further purified by size exclusion chromatography using a GE HiLoad 16/60 Superdex 200 PG (GE Healthcare Life Sciences) column connected to the AKTApurifier 100 system. A 10 mM Tris-HCl pH 8.0, 150 mM NaCl buffer was used for elution.

### Gp5β-gp5.4 complex crystallization, X-ray data collection, structure solution and refinement

The protein complex was brought to a concentration of 20 mg/ml in 10 mM Tris-HCl pH 8.0, 150 mM NaCl buffer. Initial crystallization screening was carried out employing the method of sitting drop in 96 well MRC 2 plates (SWISSCI) using Jena Bioscience crystallization screens with the help of Mosquito crystallization robot (SPT Labtech). Optimization of crystallization conditions was carried out in 24 well plates (Corning) employing the method of hanging drop vapor diffusion. Crystals were obtained by mixing 1.25 μl of purified protein complex with 1.25 μl of reservoir solution and allowed to equilibrate against 500 μl of 26% PEG 2000, 80 mM MgCl_2_, 100 mM Tris-HCl pH 8.5 at 18 °C. Crystals appeared in about 3 days quickly reaching their maximum dimensions of 0.6 mm *x* 0.1 mm *x* 0.05 mm forming quartz-like clusters. For data collection, the crystals were dipped for 20-30 seconds into the cryoprotectant solution containing 25% v/v of ethylene glycol in additional to the crystallization solution components and flash frozen in a vaporized nitrogen stream at 100° K. Crystals belonged to P2_1_, #4 space group with *a* = 46.30, *b* = 49.33, *c* = 84.06 Å, *β* = 96.19° unit cell parameters. Data collection and fluorescence scan were carried out at the X06SA PXI Pilatus beam line of the Swiss Light Source (SLS) at the Paul Scherrer Institute (SLS, Villigen, Switzerland) at the wavelength of 1 Å. Best crystals diffracted to better than 1.2 Å resolution. The diffraction data was indexed, integrated and scaled with XDS [52], details are summarized in **Supplementary Table 3**. The structure of the gp5β-gp5.4 complex was determined by molecular replacement using the program PHASER [53]. The 1.3 Å structure of the gp5R484 deletion mutant was used as a search model. Structure solution revealed that the complex of interest consists of a gp5G484 trimer and a monomer of gp5.4. The gp5.4 model was built *de novo* manually with Coot [54] and refined with SHELX L (https://doi.org/10.1107/S2053229614024218). The structure was deposited in the Protein Data Bank under the 4KU0 accession code.

### Cryo-EM

Aliquots of 3 μl of purified RB43 and T4 *5.4am* samples were applied onto QuantiFoil R1.2/1.3 Cu 300-mesh holey carbon grids (Quantifoil Micro Tools GmbH), and plunge frozen in liquid ethane with a Thermo Fisher Scientific (TFS) Vitrobot Mark III (10°С, 100% relative humidity, 3 seconds blotting time). Single particle cryo-EM data were collected at the Interdisciplinary Centre for Electron Microscopy (CIME, EPFL) in automated manner using SerialEM [55] on a FEG Tecnai F20 TEM operating at 200 kV, equipped with an Eagle 4K CCD camera (TFS).

The data were collected at the nominal magnification of 50’000, corresponding to the calibrated pixel size of 2.26 Å, with the nominal defocus range of -1.0 to -2.5 μm, and the total dose of 20 electrons per square angstrom (e^-^/Å^2^) for each exposure. A total number of 154 and 214 micrographs of RB43 and T4 *5.4am* phage particles were collected within a one-day session and imported into cryoSPARC v4.7 [56]. Contrast transfer function (CTF) was estimated with Patch CTF Estimation. Initial particle picking was performed with Blob Picker (circular and elliptical blobs of 400–900 Å diameters, a minimum relative separation distance of 0.5). In total, 96’506 RB43 particles and 155’255 *T4 5.4am*particles and were picked and extracted with a box size of 512 pixels, Fourier cropped to 128 pixels (binning 4 times) and subjected to the 2 rounds of reference-free 2D classification.

The best 1’637 RB43 baseplate particles and 1’904 *T4 5.4am* baseplate particles were selected, subjected to Ab-Initio Reconstruction with 2 Ab-Initio classes, and further refined with Non-Uniform Refinement and Local Refinement with C6 symmetry imposed. Resolution was measured with Validation (FSC) of 17.7Å for RB43 baseplate and of 17.5Å for *T4 5.4am* baseplate. Cryo-EM reconstructions were colored by cylinder radius with a rainbow palette, and figures were prepared with UCSF Chimera [57].

Reconstructions will be provided upon request.

### P2 gpV localization assay

P2 Vir1 V*am* was produced as described previously [37]. Briefly, 1 L of C-520 F+ *supD E. coli* strain at A_600_ = 0.1 was infected with one plaque of P2 Vir1 V*am*, supplemented with 0.5 mM CaCl_2_, 1.5 mM MgCl_2_, 0.1 % glucose, and grown to lysis when phage reabsorption was stopped by the addition of 0.15% EDTA. The phage was precipitated from this mixture by the addition of PEG 6000 to a final concentration of 8% w/v and NaCl to a concentration of 2.5% w/v. The phage pellet was dissolved in the SM buffer without gelatin (50 mM Tris-HCl pH 7.5, 8 mM MgSO_4_, 100 mM NaCl) and the titer was estimated by a standard plaque assay. Pure phage stocks were produced by isopycnic centrifugation at 50,000 x g for 12 hours at 4 °C in the SM buffer supplemented by CsCl to achieve an average density of 1.45 mg/ml.

For gpV localization, 100 A_600_ units of C-2 *E. coli* (∼10^11^ cells) were mixed with 5×10^12^ pfu of P2 Vir1 V*am* or an equivalent volume of SM buffer for noninfected control and incubated for 15 min at 37 °C. The cells were pelleted by centrifugation for 5 min at 5,000 *x g*. To prepare the total soluble (TS in **Fig. 4**) and total insoluble (TI) fractions (Pathway 1 in **Fig. 4**), the cell pellet (CP) was resuspended in 10 ml of buffer S (10 mM Tris-HCl, pH 7.5), the bacteria were lysed by sonication, and the mixture was centrifuged at 100,000 *x g* for 1.5 h at 4 °C. To prepare the soluble periplasmic (SP), soluble cytoplasmic (SC), and membranes and other insoluble (M) fractions (Pathway 2 in **Fig. 4**), the cell pellets were resuspended in 1 ml of sucrose-EDTA-lysozyme buffer (200 mM Tris-HCL, pH 8.0, 500 mM sucrose, 1 mM EDTA, 1 mg/mL lysozyme), incubated on ice for 30 min and centrifuged at 16,000 *x g* for 30 minutes at 4 °C. The supernatant is the SP fraction. The pellet was resuspended in 10 ml of buffer S, lysed by sonication, and centrifuged at 100,000 x g for 1.5 h at 4 °C. The supernatant is the SC fraction and the remaining pellet the M fraction. For electrophoretic assays and qPCR, the CP and M pellets were resuspended in 1 ml of buffer S.

Western blotting was performed by following a standard procedure. Protein samples were first separated on a 12% SDS polyacrylamide gel and transferred to a PVDF membrane using the iBlot transfer system (Thermo Scientific). The primary antibodies used were: MalE/MBP-probe antibody (Santa Cruz Biotechnology, sc-13564), GroEL polyclonal antibody (Enzo Life Sciences, ADI-SPS-875-D), OmpF antibody (orb308741, Biorbyt) and a custom made anti-P2 gpV polyclonal antibody generated by GenScript. The secondary antibodies were a goat polyclonal antibody to rabbit IgG (ab6721) and a goat polyclonal antibody to mouse IgG (ab6789), both from Abcam. The immuno-reactive bands were detected on a UVP ChemStudio touch Imaging System (Analytik Jena) using the Amersham ECL Western blotting detection reagents (GE Healthcare) or SuperSignal West Femto for gpV detection (Thermo Scientific).

qPCR was performed using ThermoScientific Maxima SYBR Green/ROX qPCR Master Mix. The following pair of primers was used CGCGCAGAGTTTCGGAAAAA-TCAACCTTTCGCCCCGTAAA. The qPCR reaction amplified the region 30,385-30,482 of the P2 genome that codes for orf91, an essential protein. Calibration curves were used to transfer Ct value into PFU value. Serial 10-fold dilutions of a CsCl-purified stock of P2 Vir1 V*am* with a known PFU value were used to make the calibration. Fractions with high concentration of material (like the SC) were diluted to fit the linear section of the calibration curve.

### Phage production

Primary lysates were produced on liquid cultures of the indicated *E. coli* strains according to the standard protocol [51]. Secondary lysates were prepared in supplemented LB low salt (10 g l-1 tryptone, 5 g l-1 yeast extract, 5 g l^-1^ NaCl, 2.5 g l^-1^ NH_4_Cl, 1 mM MgSO_4_, 0.1 mM CaCl_2_, 50µg ml^-1^ tryptophan). Phages were concentrated by PEG precipitation and resuspended in a small volume of SM low salt buffer (25 mM Tris HCl, 50 mM NaCl, pH 7.5) supplemented with 10 mM MgSO_4_ (SMls+Mg). Alternatively, some secondary lysates were prepared by collecting superinfected bacteria before cell lysis [51]. Lysis was then induced by treating the cell pellet resuspended in SMls+Mg with CHCl_3_. After addition of DNAse I and RNAse A (1 µg ml-1 each), the lysates were incubated for 40 min at 30°C, and bacterial debris were removed by centrifugation at 2000 x *g* for 10 min. Phages were purified by isopycnic CsCl gradient centrifugation according to the protocol of the Center for Phage Technology (Texas A&M University) and dialyzed overnight at 4°C against three changes of SM buffer with decreasing NaCl concentration (1M, 100mM, 50mM).

### Determination of phage infection and stability parameters

For all experiments, bacteria were grown to an OD_600_ = 0.5. Phage stocks used were produced on strain B^E^ and purified on CsCl gradients, except for burst size determination, for which we used phage stocks produced on a *supF* strain. *Efficiency of plating (EOP)*: T4+ and T4 *5.4am* phage titers were divided to the respective mean phage titer determined on the wild-type control strain. *Adsorption*: phages and bacteria (MOI = 0.02) were incubated 10 min at 37° and centrifuged 2 min (6000 x *g*, 4°C). For each sample, a control experiment without bacteria was also performed in parallel. The number of pfu remaining in the supernatant (Rpfu) was determined with strain B^E^. Adsorption efficiencies (i.e. % of phage attached to bacteria) were calculated as follow: 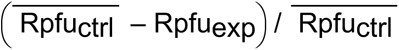.

#### Efficiency of center of infection (ECOI)

Bacteria were infected with phages (MOI = 3.5), incubated 10 min at 37°C, centrifuged twice 2 min (6000 x *g*, RT), washed with fresh growth media, and plated in top agar with strain B^E^ to determine the number of infective centers (IC). ECOI was calculated by dividing IC titer by the respective mean IC titer determined on the wild-type strain. *Burst size*: Bacteria were infected with phages (MOI = 0.01), incubated 5 min at 37°C, and centrifuged 2 min (6000 x *g*, RT). The bacterial pellet was resuspended in fresh growth media and grown with agitation at 37°C. IC at 5 and 30 min after resuspension were determined with strain B^E^. For each replicate, the number of free phages at 5 min (determined with chloroform) was subtracted from IC value and burst size was determined by dividing IC_30min_ by IC_5min_. *Stability*: Phage stocks were diluted in buffer BU (50mM Na_2_HPO_4_, 22mM KH_2_PO_4_, 70mM NaCl) at a concentration of 5*10^5^ pfu ml^-1^. Five ml of those solutions were incubated in hermetically closed plastic tubes at 37°C for the indicated time. All other T4 phage procedures were done according to standard protocols [51].

### Intracellular phage growth and phage competition assays

Intracellular phage growth assays were performed as previously described [39, 58] except that LB medium supplemented with glucose and tryptophan was used to grow bacteria. Competition assays in supplemented LB medium and determination of the ratio of *5.4*^+^ to total phage were performed as [39]. Primers SegCup (5’-AGATCCTCCTGCTCCAGTAAG) and 5.4dwn (5’-CCGACGAATATCTCGGGAATAG) were used for PCR amplification and PCR products were digested with PvuII.

#### Transposon mutagenesis and screen for T4 5.4am R T4+ S bacteria

A library of 100’000 Tet^R^ insertion mutants was generated in strain B178 by mini-Tn*10* transposon mutagenesis using the lambda 1098 phage [59]. The pooled mutants were grown to mid-log phase and 5*10^6^ bacteria were infected with T4 *5.4am* phages (produced on a *sup*^+^ strain) at a MOI of 10 in 200µl of growth media for 20 min at 37°C. The mixture was then poured in a soft HA overlay on HA-agar plate. After overnight incubation, we observed about 200 phage-resistant colonies. Thirty-two individual colonies were purified and individually tested for resistance to T4+ and T4-*5.4am*. Three of 32 were resistant to T4 *5.4am* but sensitive to T4+, all the other mutants were resistant to both wild type and mutant phages. The transposon insertion sites were mapped according to published methods [60]. The three mutant strains have a transposon inserted in gene *hldD* at the exact same site suggesting that they might derive from the same insertion event.

### In vivo and in vitro complementations

*In vivo complementations* Rosetta (DE3) bacteria (*sup^+^*) harboring plasmid pET23d or pET23d-*5.4* were grown in 3ml of LB low salt at 37°C to OD_600_ = 0.3, induced with 1mM IPTG for 30 min, infected with 3.4*10^8^ pfu of T4 *5.4am* (produced on a *sup^+^*strain), superinfected 6 min later with the same number of phages, and incubated for 1.5 h with agitation. Remaining bacteria were killed and permeabilized with CHCl_3_, and cell debris were removed by low-speed centrifugation at 4°C. *In vitro complementation* Cell extracts for *in vitro* complementations were produced from Rosetta (DE3) bacteria harboring plasmid pET23d or pET23d-*5.4* grown and induced as described above without phage infection. Induced bacteria were immediately permeabilized as described above and cleared cell lysates were kept at 4°C before use. Complementations were done by incubating for 45 min at 37°C equal volume of T4 *5.4am* phages produced on Rosetta (DE3) bacteria harboring plasmid pET23d and fresh cell extracts produced from bacteria harboring plasmid either pET23d or pET23d-*5.4*.

## Data availability

Full gel/blot images and numerical data used to generate panels and graphs in Figures 4, 5, 6, and S2 are contained in the corresponding source data files (Figure X-source data_n). Diffraction data for the crystal structure displayed in Figure 2 has been deposited in PDB under the accession code 4KU0

Full 3D cryo-EM reconstruction and corresponding FSC files of the baseplate regions of T4 5.4am and RB43 displayed in Figure 3 and S1 have been stored in cloud storage allowing data upload upon request.

The genome sequence of T4-5.4am has been deposited in GenBank under the accession code PX867600

## Author Contributions

Conceptualization, Y.M. E.K. P.L. & D.B; biological investigations, Y.M. E.K.; structure determination, M.M.S, S.B., S.N. & P.L.; NGS and genome analysis, W.R.; writing, Y.M. P.L. & D.B; all authors have read and agreed to this version of the manuscript.

## Acknowledgements

Y.M. & D.B. thank Costa Georgopoulos for many helpful discussions and Filo Silva for technical help with cloning, transposon mutant screening and competition experiments. D.B. thanks the late Dick Epstein for discussions about the T4 ORFans. Y.M. thanks Patrick Viollier for supporting his research on phages during the last years.

