## Supplementary Material for "The spike tip protein of bacteriophage T4"

#### Supplementary Materials and Methods

##### Plasmid construction

pET23d_*5.4* was constructed by PCR amplification of gene 5.4 using the following primer pair, 5.4F 5'-ATACCATGGCAGGATTAAGTTATGATAAGTGT / 5.4R 5'-ATAAAGCTTATTTAATGAATACTTTAGATGATG, and cloning in the NcoI and HindIII site of pET23d. This vector expresses the native protein (without a His-tag), because a stop codon is included in the reverse primer (underlined). pBR322ΔTc was constructed by digesting pBR322 with EcoRV and NruI, followed by self-ligation.

##### Bacterial strain construction

To construct YM66, the *ΔompC*::*kan* allele of JW2203 was converted to a *ompC*::*frt* allele by removing the *kan* cassette with FLP recombinase supplied from plasmid pCP20 as described [1]. To construct YM67, a P1 lysate was prepared following standard procedure [2] on strain JW3596 (*ΔrfaC*::*kan* ) harboring plasmid pCA24N-*rfaC* in presence of 1mM IPTG and it was used to transduce the *ΔrfaC*::*kan* allele in strain YM66.

##### T4 strain construction.

A *5.4* amber mutant of the Tyr6 codon (TAT -> TAG mutation) was constructed using the T4 I/S system [3] [4]. This mutation does not affect the end of the *segC/5.3* gene that overlaps the first 5 codons of gene *5.4*. To facilitate the detection of *5.4am* mutant phage, a silent mutation that generates a PvuII restriction site was introduced at the Thr11 codon (ACT -> ACA). We constructed by PCR site-directed mutagenesis [5] a 590 bp fragment containing the last 321 bp of the *segC/5.3* gene, the first 252 bp of gene *5.4* gene with the above-described mutations, and 15bp homology sequences to the multiple cloning site of pBSPL0+ on each side of the fragment (external primers: 5.4_supf_mut1 5'-AGCTTGTCGACTCGAGGCGGTATCGGTGGCT / 5.4_supf_mut2 5'-GTACCATATGCTCGA GCCGCATGATATTGGATCA, 5.4_supf_mut3 5'-CAGGATTAAGTTAGGATAAGTGTGTT ACAGCTGGCCATG / 5.4_supf_mut4 5'-CATGGCCAGCTGTAACACACTTATCCTAA CTTAATCCTG, mutations underlined). The fragment was cloned in pBSPL0+ linearized with XhoI using the InFusion kit (Clontech/Takara Bio) and the insert was sequenced. The were then transferred to the T4 K10 genome. The presence of the *5.4am* and PvuII mutations was confirmed by PCR amplification of the *segC-5.4* region, followed by PvuII digest and sequencing. The *5.4am* mutation was transferred from the T4 K10 *5.4am* to a clean T4D genetic background in two steps. First, T4 K10-*5.4am* was crossed with T4D (MOI of 5 for each phage) and the progeny was plated on B178(λ) (a strain that carries an uncharacterized amber suppressor) to select against the rIIPT8 mutation (*denB-rII* deletion) present in the K10 background. Individual plaques were screened as described above to identify *5.4am* mutants. Secondly, one of these *5.4am* $denB^+$ $rII^+$ phage was crossed with T4D and the progeny was plated on $B^E$ to select against the *38am* and *51am* mutations. The *5.4am* and $denA^+$ genotype of the chosen mutant was confirmed by PCR and sequencing.

The T4-K10 strain was shown to carry a few mutations in addition to the four mutations described above [6]. Therefore, the genome of the *5.4am* strain used in this study was fully sequenced. DNA was extracted from purified intact phage particles and then used to generate Illumina NGS libraries for single-end sequencing on an Illumina HiSeq 2500 system. As T4D, the wild-type T4 strain used in this study, has not been sequenced, approximately 7.7 million 50 nucleotide reads were mapped to the T4T genome (Accession HM137666.1) [7]. This reference was selected because of the high degree of nucleotide identity when compared to other available T4 isolated genomes. Mapping, mutational confirmation, and SNP identification was completed using CLC Genomics Workbench v8 (CLC Genomics). SNPs with more than 99% coverage and read depth of 600 were identified as potential suppressors. In addition to the *5.4am* and PvuII mutations, thirteen SNP were identified (see table S2). Three of these are synonymous mutations while at four other loci the predicted protein sequences encoded by the *T4-5.4am* phage match most corresponding T4 and T4-like published protein sequences. Of the remaining six SNP, one is found in the conserved RNA gene, *rnaD*, and all the other are missense mutations (*dda*, *e.8*, *30*, *34*, and *ndd*). We have PCR-amplified and sequenced these loci in the wild-type T4D strain used in this study and found that it harbors the same mutations as T4-*5.4am* in genes *dda*, *30*, *34*, and *ndd*. Surprisingly the mutations in genes *rnaD* and *e.8* are present neither in T4D nor in T4-K10. Thus, they must have arisen during the construction of the T4 *5.4am* strain.

**Table S1.** List of mutations in T4 *5.4am* compared to the reference T4T sequence

| | T4T | | T4-*5.4am* | T4D | T4T→ T4-*5.4am* | T4-like |
|---|---|---|---|---|---|---|
| **Gene** | **Position** | **Base (+ strand)** | **Base** | **Base** | **aa change** | **Conserved aa** |
| *rIIA* | 722 | T | C | nt | T490→A | A |
| *motB.1* | 7421 | G | A | nt | syn | |
| *dda* | 9542 | C | T | T | A392→T | A, V, C |
| *segF* | 15683 | T | C | nt | D195→G | G, A |
| *b-gt* | 24419 | T | C | nt | Stop341→W | W |
| *nrdC.11* | 56175 | A | G | nt | syn | |
| *e.8* | 70530 | C | A | C | D25→Y | D |
| *rnaD* | 71105 | G | A | G | na | na |
| ***5.4*** | **80775** | **T** | **G** | **T** | **Y6→stop** | **Y** |
| *5.4* | 80790 | T | A | T | syn | |
| *24* | 107807 | C | A | nt | syn | |
| *hoc* | 110431 | T | G | nt | S300→R | R |
| *30* | 127060 | C | T | T | A56→T | A |
| *34* | 151972 | C | T | T | P384→L | P |
| *ndd* | 165942 | G | C | C | P38→ R | P |

T4T, sequence according to GeneBank (HM137666.1). T4 *5.4am*, determined by full genome sequencing. T4D, determined by PCR amplification and Sanger sequencing on wild-type T4D laboratory stock. T4T→T4-*5.4am*, predicted amino acid changes between T4T and T4-*5.4am.* T4-like Conserved aa, amino-acid present at this position in homologous proteins of sequenced T4-like phages. The 5,4 amber mutation is shown in bold. na: not applicable; nt: not tested; syn: synonymous.

**Table S2.** Bacteria, phage and plasmids

| Strain | Genotype | Reference |
|---|---|---|
| **Bacterial strains** | | |
| B178 | *E. coli* K-12, W3110 *galE* | [8] |
| B178 ($\lambda$) | B178 $\lambda$ lysogen, harbors an uncharacterized amber suppressor | C. Georgopoulos |
| $B^E$ | Prototrophic, an *E. coli* B strain | [9] |
| Rosetta(DE3) | *E. coli* B, $F^-$ *ompT hsdS*$_B$ *gal dcm lacY1 λ*(DE3) pRARE (encodes the tRNA genes *argU, araW, ileX, glyT, leuW, proL, metT, thrT, tyrU and thrU*, $Cm^r$) | Novagen |
| BW25113 | *E. coli* K-12, *F- Δ(araD-araB)567 ΔlacZ4787(::rrnB-3) rph-1 Δ(rhaD-rhaB)568 hsdR514* | CGSC |
| CR63 | *E. coli* K-12, *F+, supD (serU60), lamB63* | lab collection/CGSC |
| C-2 | *E. coli* C $F^+$ *sup*$^+$ | [10] |
| C-520 | *E. coli* C $F^+$ *supD* | [10] |
| JW3596 | BW25113 *ΔrfaC733::kan* | [11] |
| JW3594 | BW25113 *ΔhldD731::kan* | [11] |
| JW3595 | BW25113 *ΔrfaF732::kan* | [11] |
| JW3597 | BW25113 *ΔrfaL734::kan* | [11] |
| JW2203 | BW25113 *ΔompC768::kan* | [11] |
| NM538 | *supF (tyrT58), hsdR514, trpR58* | [12] |
| W3110 | *F- IN(rrnD-rrnE)1 rph-1* | CGSC |
| YM65 | B178 *hldD::Tn10* | this study |
| YM66 | BW25113 *ΔompC::frt* | this study |
| YM67 | BW25113 *ΔompC::frt ΔrfaC733::kan* | this study |
| **Phage** | | |
| T4+ | T4D | [13] |
| T4 K10 | *38amB262 51amS29 nd28 (denA) rIIPT8 (ΔdenB-rII)* | [4] |
| T4 K10 *5.4am* | K10 *5.4am* (TAT -> TAG) | this study |
| T4 *5.4am* | T4D *5.4am* | this study |
| P2 *vir* V*am* | | [10] |
| **Plasmids** | | |
| pBR322 ΔTc | $Ap^r$ $Tc^s$ | this study |
| pBSPI0+ | Vector with pBR322 replication origin, expressing *supF* from $P_{23}$ (phage T4 late promoter), $Ap^r$ | Selick et al. 1988 |
| pCA24N(-) | Vector with pBR322 replication origin, $P_{T5\text{-}lac}$ (IPTG-inducible T5 promoter), *lacI*$^q$, $Cm^r$ | [14] |
| pCA24N-*hldD* | pCA24N(-) expressing His-*hldD* from $P_{T5\text{-}lac}$ | [14] |
| pCA24N-*rfaC* | pCA24N(-) expressing His-*rfaC* from $P_{T5\text{-}lac}$ | [14] |
| pET23d | Vector with pBR322 replication origin, $P_{T7}$, $Ap^r$ | Novagen |
| pET23d_*5.4* | pET23d expressing *5.4* from $P_{T7}$ | This study |

**Table S3.** X-ray data collection and refinement statistics of gp5β-gp5.4 complex.

Data in parenthesis represent statistics for the highest resolution shell.

| | |
|---|---|
| **Data collection** | |
| Wavelength | 1.0 Å |
| Number of frames | 1440 |
| Frame width (°) | 0.25 |
| Space group | P1 2(1) 1 (#4) |
| Unit cell parameters: | |
| a, b, c (Å) | 46.30, 49.33, 84.06 |
| α, β, γ (°) | 90, 96.19, 90 |
| Biologic assembly | Tetramer (trimer + monomer) |
| Number of biological assemblies per asymmetric unit | 1 |
| Resolution (Å) | 46.0 – 1.15 |
| Rmeas | 0.057 (0.436) |
| <I / σI> | 12.1 (2.3) |
| Completeness (%) | 95.2 (81.3) |
| Redundancy | 3.2 (2.5) |
| **Refinement** | |
| Number of reflections | |
| Working | 124490 |
| Test | 4516 |
| Rwork / Rfree | 0.127/0.171 |
| B-factor (Å2) | 22.9 |
| R.m.s. deviations | |
| Bond lengths (Å) | 0.013 |
| Bond angles (°) | 0.030 |
| Number of atoms | |
| Protein | 2893 |
| Solvent and ligands | 638 |
| Ramachandran plot (%) | |
| Most favored | 95.2 |
| Additionally allowed | 4.5 |
| Outliers | 0.3 |

#### Supplementary figure legends

**Supplementary Figure 1. Location of spike-tip complex in RB43 phage. Sliced view of cryo-EM reconstructions of RB43 baseplates.**

The structures of the RB43 spike (orf204) and the putative RB43 spike tip protein (orf205w) are predicted by AlphaFold. Predicted complex rigid-body fitted into different cryo-EM reconstructions of RB43 baseplate after 3D classification focused on spike-tip complex.

**Supplementary Figure 2**. **Infection properties of T4+ and *T4 5.4am* on WT and LPS mutant cells.**

**A**, The EOP of T4+ and *T4 5.4am* produced on a non-permissive *sup*$^{+}$ strain was determined on non-permissive *sup*$^{+}$ *E. coli* K12 (B178 (WT)) and *E. coli* B ($B^{E}$), and on permissive *supF* (NM538) and *supD* (CR63) *E. coli* K-12. The EOP was calculated by dividing the phage titer obtained on the tested strain by the titer obtained on the *supF* strain. Data are the mean and standard deviations of three independent experiments.

**B**, The stability of T4+ and T4 *5.4am* was determined at the indicated time points after initial dilution at $2*10^{5}$ pfu/ml in low salt potassium buffer BU and incubation at 37°C. Data are the means and ranges of two measurements normalized to the titer at day zero.

**C**, The rate of adsorption of T4+ and *T4 5.4am* produced on a *sup*$^{+}$ strain were determined on *E. coli* $B^{E}$ (see *Material and Methods*). The titers of phage left in solution were determined at the indicated time points and normalized to the titer at time zero (100%). A representative of two independent experiments with similar outcomes is shown.

**D,** To confirm the deep-rough phenotype of the *hldD* mutant, overnight cultures of B178(WT) and *hldD::*Tn*10* were diluted and spotted on LB agar medium supplemented with the indicated chemicals and on McConkey agar.

**E**, Growth of T4+ and *T4 5.4am* on *E. coli* deletion mutants of *ompC* and of the core LPS synthesis genes. Phage stocks produced on the *sup*$^{+}$ strain $B^{E}$ were diluted and spotted onto bacterial lawns of WT (BW25113), *ΔhldD* (JW3594-1), *ΔrfaF* (JW3595-2), *ΔrfaC* (JW3596-1), *ΔwaaL* (JW3597-1), *ΔompC* (JW2203-1), or *ΔompC ΔrfaC*

(YM65) (see Supplementary Table 2 for strain references). Results are representative of three independent experiments.

### Supplementary Figure 1

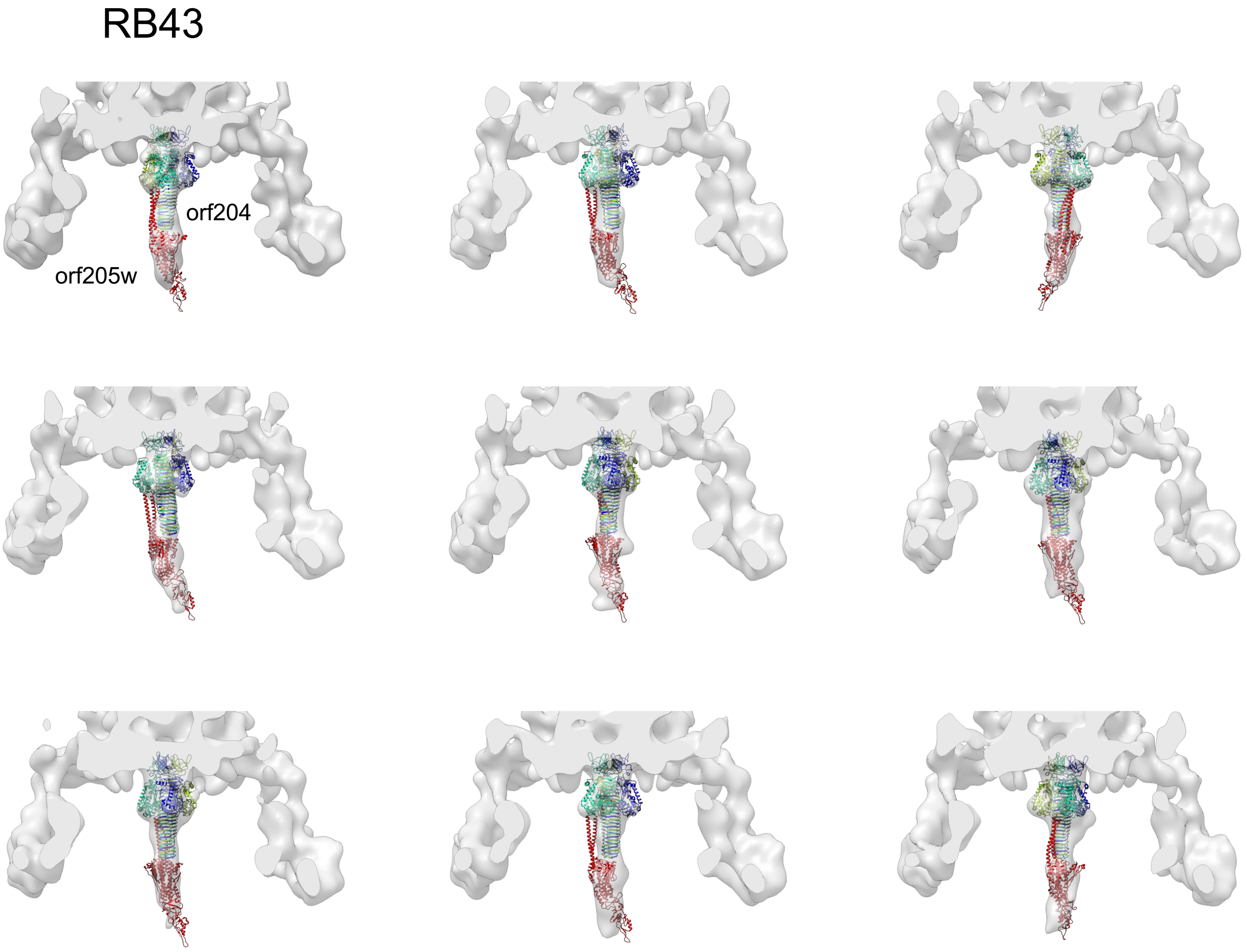

### Supplementary Figure 2

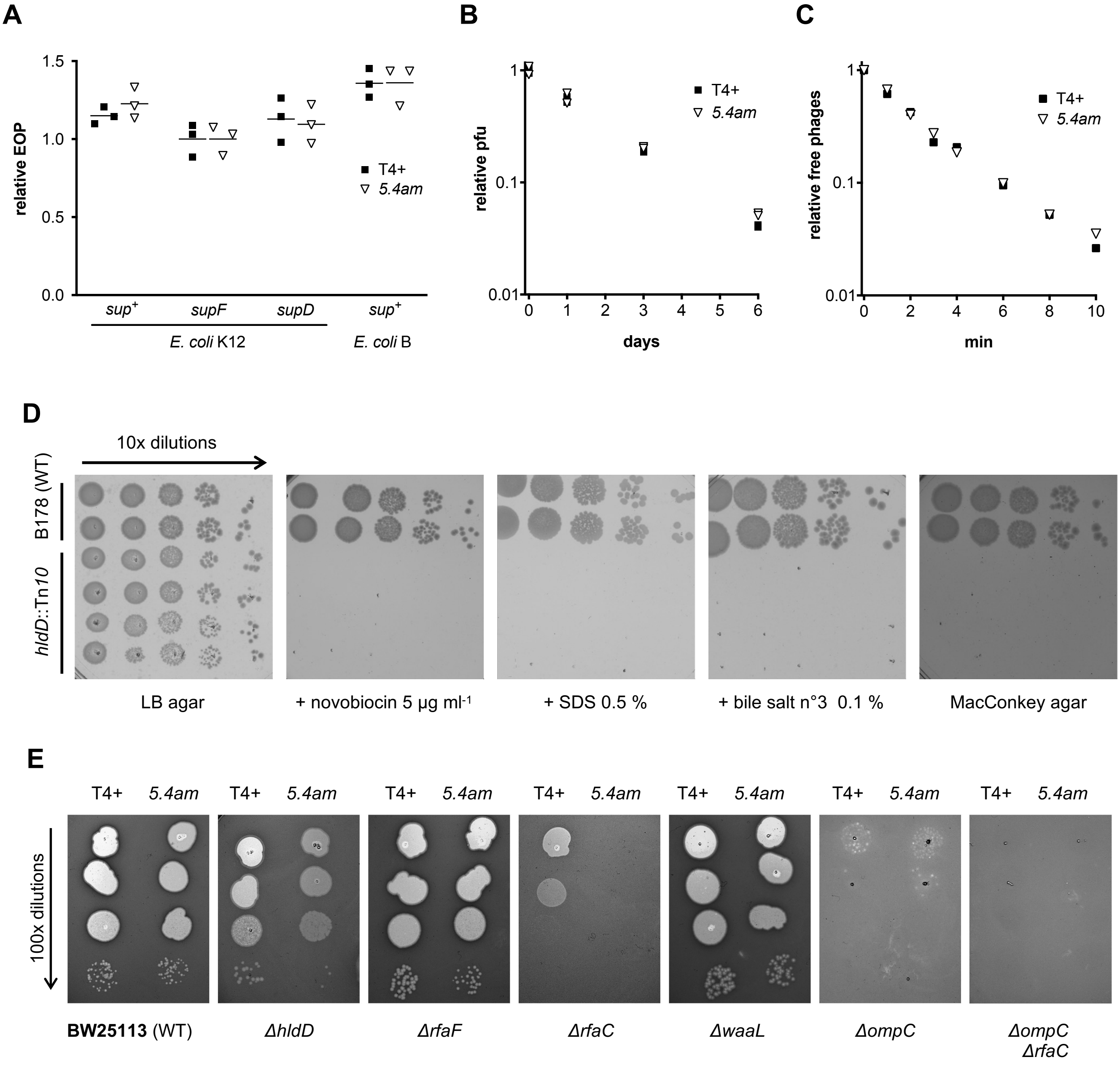

#### Figure 4-source data_panel B

| replicate | Initial titer (%) | Titer after absorbption (%) | Initial titer (%) | Titer after absorbption (%) |
|---|---|---|---|---|
| 1 | 100 | 5 | Average | Average |
| 2 | 100 | 1.4 | 100 | 4.13333 |
| 3 | 100 | 6 | StDev | StDev |
| | | | 0 | 2.41937 |

### Figure 4-source data_panel C

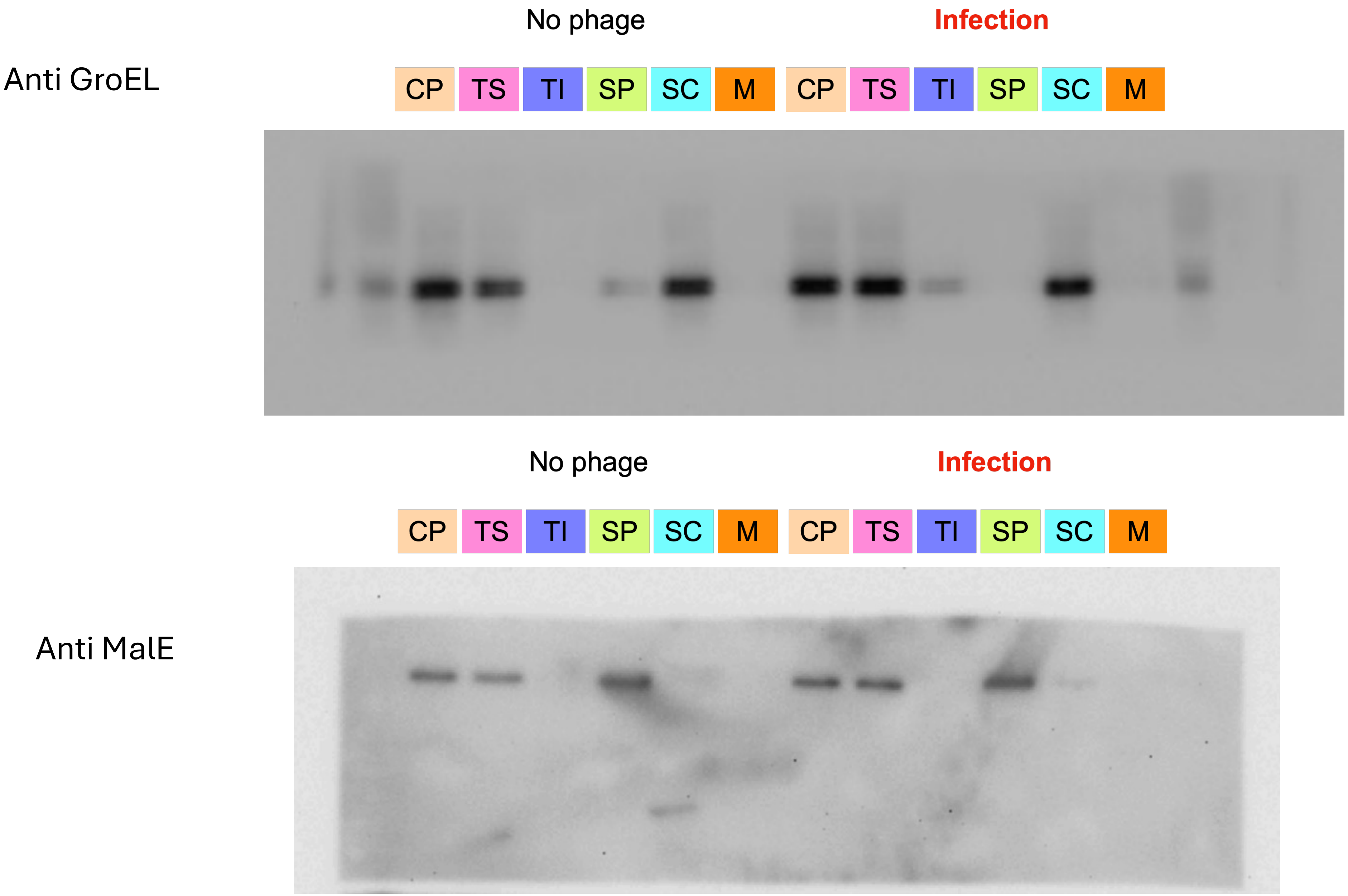

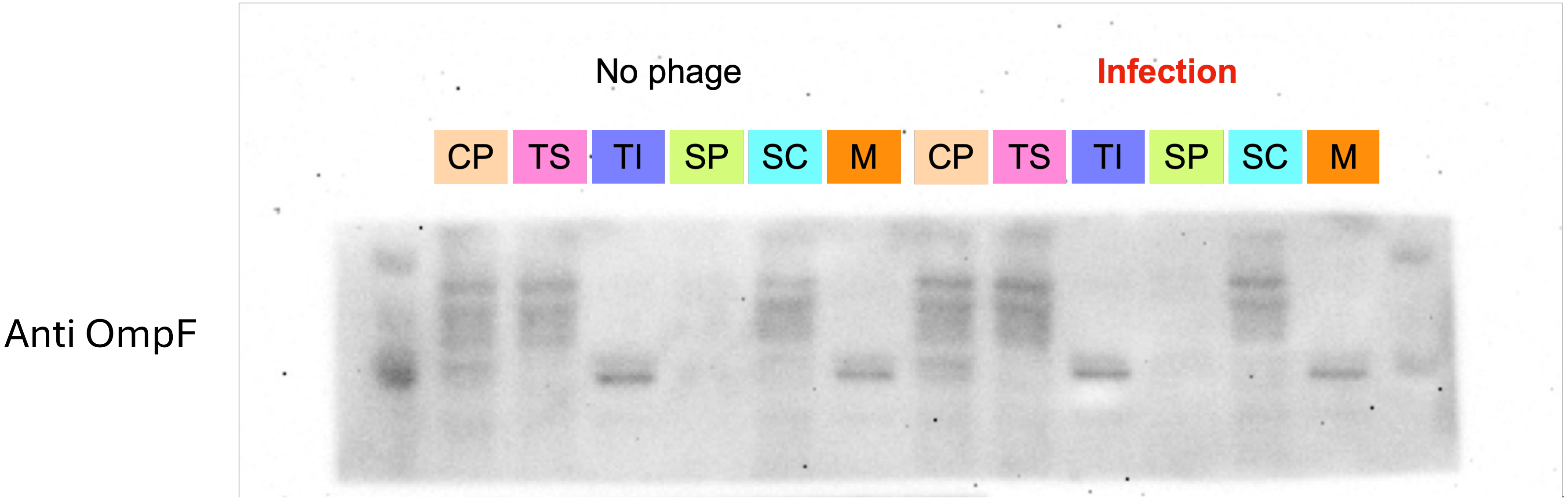
No phage
Infection
CP
TS
TI
SP
SC
M
CP
TS
TI
SP
SC
M
Anti OmpF

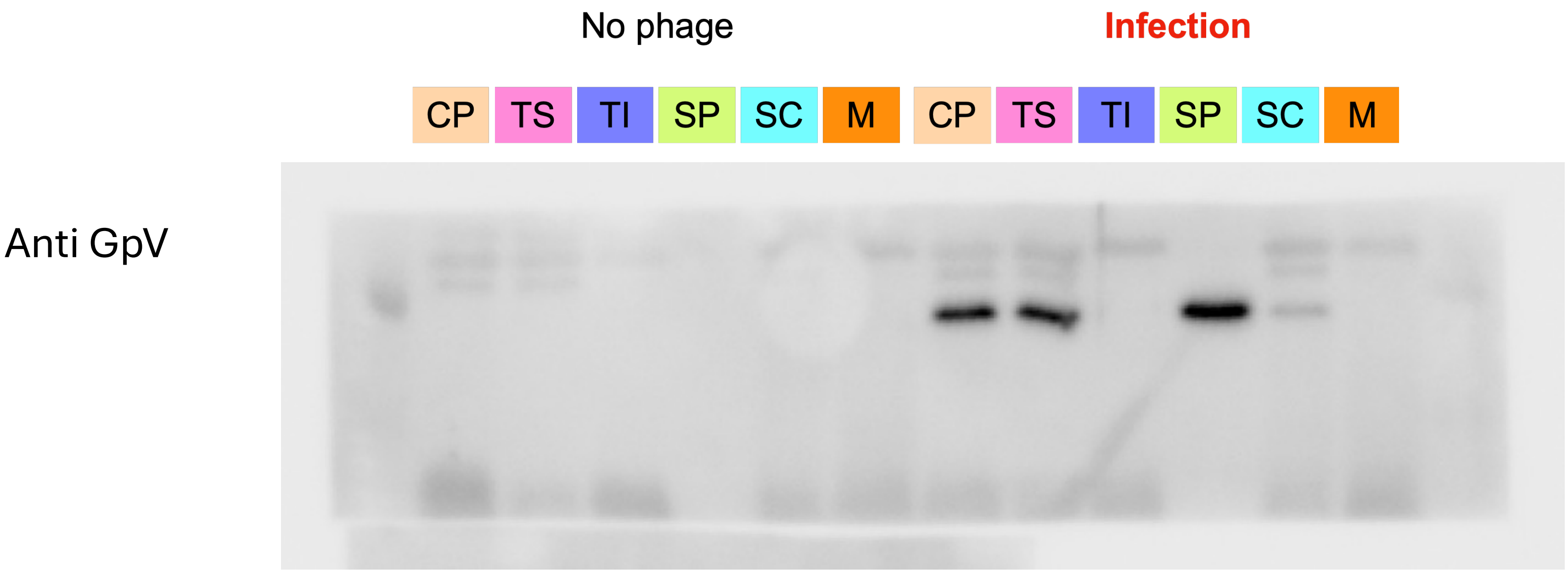
No phage
Infection
CP
TS
TI
SP
SC
M
CP
TS
TI
SP
SC
M
Anti GpV

### Figure 4-source data_panel D

Calibration curve

| pfu | log10(pfu) | Ct rep. 1 | Ct rep. 2 | Ct rep. 3 | Av. Ct |
|---|---|---|---|---|---|
| 9.00E+07 | 7.9542 | 6.956678 | 6.892436 | 6.939355 | 6.929490 |
| 9.00E+06 | 6.9542 | 9.231696 | 9.586086 | 9.712253 | 9.510012 |
| 9.00E+05 | 5.9542 | 12.508165 | 12.727257 | 12.632950 | 12.622791 |
| 9.00E+04 | 4.9542 | 16.509005 | 16.317144 | 16.460171 | 16.428773 |
| 9.00E+03 | 3.9542 | 19.893459 | 19.819715 | 19.901260 | 19.871478 |
| 9.00E+02 | 2.9542 | 23.177387 | 23.451702 | 23.234720 | 23.287937 |
| 9.00E+01 | 1.9542 | 26.802420 | 26.710419 | 26.605633 | 26.706157 |
| 9.00E+00 | 0.9542 | 28.709263 | 28.553354 | 27.877106 | 28.379908 |

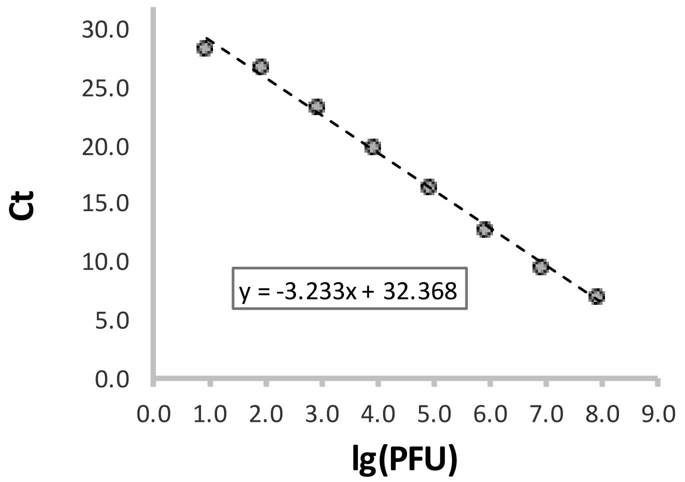

| | Ct rep. 1 | Ct rep. 2 | Ct rep. 3 | Av. Ct | Calibrated | pfu before dilution/vol. adjustments | By dilution factor | By dilution factor | pfu in fraction (adjusted to fraction | Log10(pfu) |
|---|---|---|---|---|---|---|---|---|---|---|
| Tot bact | 12.311872 | 12.230824 | 12.333426 | 12.292041 | 6.209700 | 1620689.5 | | | | |
| CP | 12.339796 | 12.147804 | 12.316186 | 12.267929 | 6.217158 | 1648761.4 | 164876144.3 | 54958714.8 | 3.627E+12 | 12.5596 |
| SP | 8.949521 | 9.011724 | 9.041590 | 9.000945 | 7.227669 | 16891546.7 | 168915466.7 | 56305155.6 | 2.815E+10 | 10.4495 |
| SC | 13.287230 | 13.695021 | 13.513778 | 13.498676 | 5.836475 | 686238.4 | 68623838.6 | 22874612.9 | 1.830E+11 | 11.2624 |
| M | 15.879118 | 15.795842 | 15.795503 | 15.823488 | 5.117387 | 131034.9 | 1310349.3 | 436783.1 | 4.368E+08 | 8.6403 |

### Figure 5-source data_panel A

**Single growth cycle of T4+ and T4 *5.4am***

| | T4+ | | | | | | 5.4*am* | | | | | |
|---|---|---|---|---|---|---|---|---|---|---|---|---|
| t (min) | biol. repli. | dilution $10^x$ | vol (µl) | pfu count | pfu/ml | rel. pfu/ml* | biol. repli. | dilution $10^x$ | vol (µl) | pfu count | pfu/ml | rel. pfu/ml* |
| 8 | 1 | -1 | 200 | 30 | 1.50E+03 | 3.57E-02 | 1 | -1 | 200 | 32 | 1.60E+03 | 3.76E-02 |
| | 2 | -1 | 200 | 29 | 1.45E+03 | 3.45E-02 | 2 | -1 | 200 | 29 | 1.45E+03 | 3.41E-02 |
| 11 | 1 | -1 | 100 | 22 | 2.20E+03 | 5.24E-02 | 1 | -1 | 100 | 7 | 7.00E+02 | 1.65E-02 |
| | 2 | -1 | 100 | 9 | 9.00E+02 | 2.14E-02 | 2 | -1 | 100 | 9 | 9.00E+02 | 2.12E-02 |
| 14 | 1 | -1 | 150 | 59 | 3.93E+03 | 9.37E-02 | 1 | -1 | 150 | 79 | 5.27E+03 | 1.24E-01 |
| | 2 | -1 | 150 | 80 | 5.33E+03 | 1.27E-01 | 2 | -1 | 150 | 65 | 4.33E+03 | 1.02E-01 |
| 17 | 1 | -2 | 50 | 289 | 5.78E+05 | 1.38E+01 | 1 | -2 | 50 | 287 | 5.74E+05 | 1.35E+01 |
| | 2 | -2 | 50 | 285 | 5.70E+05 | 1.36E+01 | 2 | -2 | 50 | 197 | 3.94E+05 | 9.27E+00 |
| 20 | 1 | -3 | 100 | 212 | 2.12E+06 | 5.05E+01 | 1 | -3 | 100 | 211 | 2.11E+06 | 4.96E+01 |
| | 2 | -3 | 100 | 279 | 2.79E+06 | 6.64E+01 | 2 | -3 | 100 | 180 | 1.80E+06 | 4.24E+01 |
| 23 | 1 | -3 | 75 | 303 | 4.04E+06 | 9.62E+01 | 1 | -3 | 75 | 325 | 4.33E+06 | 1.02E+02 |
| | 2 | -3 | 75 | 355 | 4.73E+06 | 1.13E+02 | 2 | -3 | 75 | 246 | 3.28E+06 | 7.72E+01 |
| 26 | 1 | -3 | 50 | 191 | 3.82E+06 | 9.10E+01 | 1 | -3 | 50 | 162 | 3.24E+06 | 7.62E+01 |
| | 2 | -3 | 50 | 173 | 3.46E+06 | 8.24E+01 | 2 | -3 | 50 | 132 | 2.64E+06 | 6.21E+01 |

* relative to phage input

| phage input | pfu/ml |
|---|---|
| T4+ | 4.20E+04 |
| 5.4*am* | 4.25E+04 |

### Figure 5-source data_panel B

**T4+ ratio (%)**

| | supD (CR63) | | sup+ (BE) | | trend | trend |
|---|---|---|---|---|---|---|
| growth cycles | biol rep1 | biol rep2 | biol rep1 | biol rep2 | simul. BE | simul. CR63 |
| 0 | 8.87390 | 7.45041 | excl. | 8.26744 | 8.00000 | 8.00000 |
| 1 | 10.00350 | 9.71453 | 8.28028 | excl. | 11.87739 | 9.05496 |
| 2 | 12.92373 | 14.13491 | 10.97491 | 18.53020 | 17.28106 | 10.23357 |
| 3 | 14.27008 | 13.26002 | 23.80443 | 23.66195 | 24.46076 | 11.54611 |
| 4 | 15.58081 | 15.16303 | 25.41071 | 34.37121 | 33.41827 | 13.00260 |
| 5 | 17.93723 | 12.84218 | 40.24561 | 45.23716 | 43.75596 | 14.61248 |
| 6 | 18.39108 | 15.88468 | 53.12192 | 58.30352 | 54.66595 | 16.38414 |
| 7 | 19.48641 | 16.32435 | 61.66587 | 65.61126 | 65.14542 | 18.32451 |
| 8 | 20.34472 | 20.39013 | 76.76288 | 74.52980 | 74.33955 | 20.43850 |

CR63 rep 1

| Growth cycle | band | QL-BG | WT percent |
|---|---|---|---|
| 0 | 1 | 2502361.19 | 8.87390 |
| | 2 | 17510820.2 | |
| | 3 | 8185930.25 | |
| 1 | 1 | 2233441.81 | 10.00350 |
| | 2 | 13136096.5 | |
| | 3 | 6957065.5 | |
| 2 | 1 | 3103999.91 | 12.92373 |
| | 2 | 13486739 | |
| | 3 | 7427092.38 | |
| 3 | 1 | 3979911.33 | 14.27008 |
| | 2 | 15149655 | |
| | 3 | 8760337.15 | |
| 4 | 1 | 4676018.24 | 15.58081 |
| | 2 | 16118618.4 | |
| | 3 | 9216759.06 | |
| 5 | 1 | 5328468.41 | 17.93723 |
| | 2 | 15459550.6 | |
| | 3 | 8918168.84 | |
| 6 | 1 | 5856139.06 | 18.39108 |
| | 2 | 16533176.9 | |
| | 3 | 9452956.55 | |
| 7 | 1 | 5926925.46 | 19.48641 |
| | 2 | 15611287.6 | |
| | 3 | 8877480.94 | |
| 8 | 1 | 13707570.4 | 20.34472 |
| | 2 | 33973759.4 | |
| | 3 | 19695219.6 | |

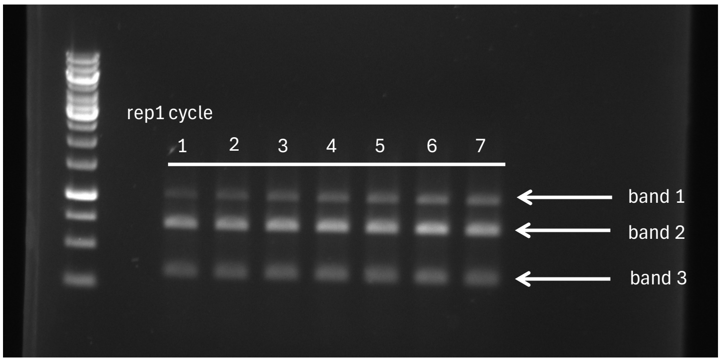

N.B. band 1 correspond to WT PCR product (undigested) and band 2 and 3 to *5.4am* PCR product (digested)

CR63 rep 2

| Growth cycle | band | QL-BG | WT percent |
|---|---|---|---|
| 0 | 1 | 605332.95 | 7.45041 |
| | 2 | 4829400.26 | |
| | 3 | 2690091.14 | |
| 1 | 1 | 6531991.73 | 9.71453 |
| | 2 | 39059564.9 | |
| | 3 | 21647819.2 | |
| 2 | 1 | 4041859.57 | 14.13491 |
| | 2 | 16557218.2 | |
| | 3 | 7995794.23 | |
| 3 | 1 | 3596877.89 | 13.26002 |
| | 2 | 15768949 | |
| | 3 | 7759915.85 | |
| 4 | 1 | 4221133.17 | 15.16303 |
| | 2 | 15787014.6 | |
| | 3 | 7830181 | |
| 5 | 1 | 3326925.29 | 12.84218 |
| | 2 | 15181133.5 | |
| | 3 | 7398174.07 | |
| 6 | 1 | 4758857.96 | 15.88468 |
| | 2 | 17440006.4 | |
| | 3 | 7759929.56 | |
| 7 | 1 | 6114010.94 | 16.32435 |
| | 2 | 21202315.6 | |
| | 3 | 10137001.4 | |
| 8 | 1 | 8705493.25 | 20.39013 |
| | 2 | 22544460.4 | |
| | 3 | 11444698.1 | |

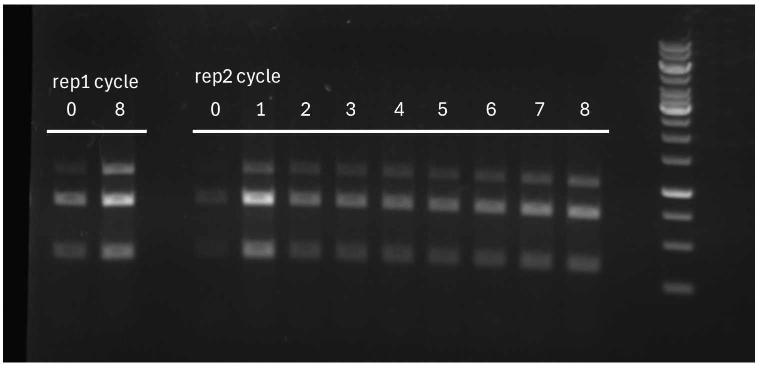

BE rep 1

| Growth cycle | band | QL-BG | WT percent | |
|---|---|---|---|---|
| *0* | *1* | *7376511.3* | *23.78352* | * |
| | *2* | *16152553* | | |
| | *3* | *7486161.7* | | |
| 1 | 1 | 408082.71 | 8.28028 | |
| | 2 | 2012388.71 | | |
| | 3 | 2507898.32 | | |
| 2 | 1 | 1224026.94 | 10.97491 | |
| | 2 | 5038376.12 | | |
| | 3 | 4890552.19 | | |
| 3 | 1 | 1042164.41 | 23.80443 | |
| | 2 | 1947538.83 | | |
| | 3 | 1388324.17 | | |
| 4 | 1 | 1903782.7 | 25.41071 | |
| | 2 | 3228366.04 | | |
| | 3 | 2359898.88 | | |
| 5 | 1 | 3227524.3 | 40.24561 | |
| | 2 | 2826248.86 | | |
| | 3 | 1965795.93 | | |
| 6 | 1 | 4756915.28 | 53.12192 | |
| | 2 | 2567360.12 | | |
| | 3 | 1630437.3 | | |
| 7 | 1 | 4863229.63 | 61.66587 | |
| | 2 | 1807532.95 | | |
| | 3 | 1215657.03 | | |
| 8 | 1 | 9152796.36 | 76.76288 | |
| | 2 | 1499381.46 | | |
| | 3 | 1271289.5 | | |

BE rep 2

| Growth cycle | band | QL-BG | WT percent | |
|---|---|---|---|---|
| 0 | 1 | 459796.71 | 8.26744 | |
| | 2 | 3457748.4 | | |
| | 3 | 1643994.43 | | |
| *1* | *1* | *9433278.8* | *68.36550* | * |
| | *2* | *2867557* | | |
| | *3* | *1497467.8* | | |
| 2 | 1 | 2388108.31 | 18.53020 | |
| | 2 | 6972631.22 | | |
| | 3 | 3526915.23 | | |
| 3 | 1 | 3233853.38 | 23.66195 | |
| | 2 | 6769782.36 | | |
| | 3 | 3663256.91 | | |
| 4 | 1 | 4960503.98 | 34.37121 | |
| | 2 | 6273986 | | |
| | 3 | 3197657.02 | | |
| 5 | 1 | 9328990.38 | 45.23716 | |
| | 2 | 7241269.6 | | |
| | 3 | 4052143 | | |
| 6 | 1 | 14398216.8 | 58.30352 | |
| | 2 | 6915457.04 | | |
| | 3 | 3381603.64 | | |
| 7 | 1 | 9294320.93 | 65.61126 | |
| | 2 | 3128627 | | |
| | 3 | 1742792.37 | | |
| 8 | 1 | 12320953.5 | 74.52980 | |
| | 2 | 2659855.05 | | |
| | 3 | 1550771.2 | | |

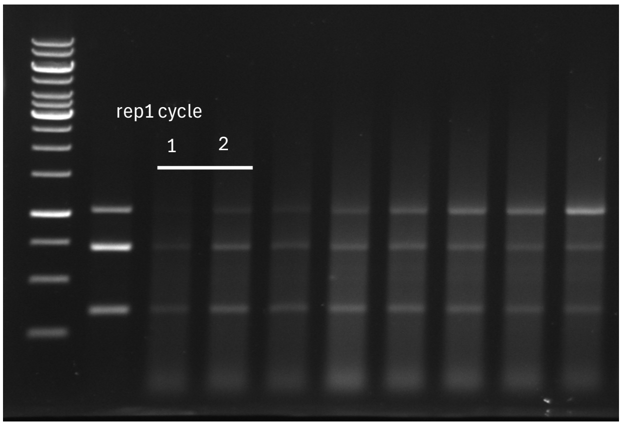

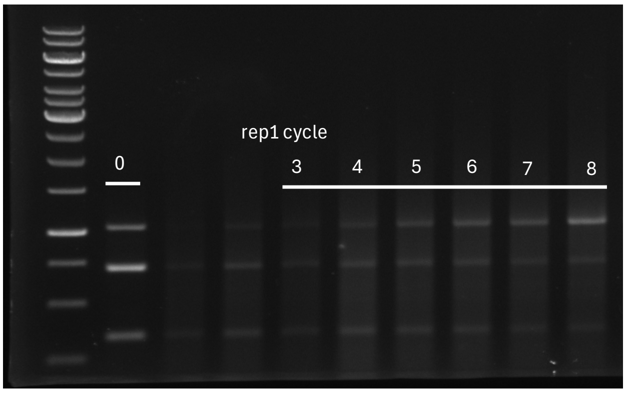

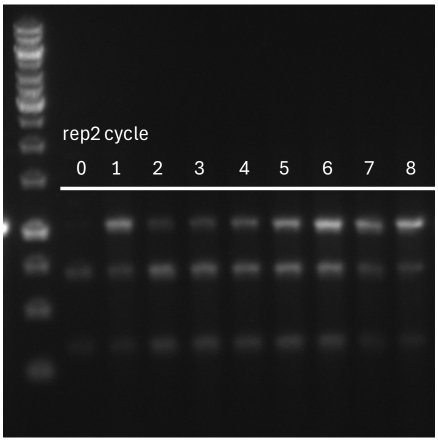

* value for rep1 growth cycle 0 and rep2 growth cycle 1 are clearly anomalous and wer excluded from analysis

### Figure 5-source data_panel C

**Infectivity**

| CsCl Sample ID | Tech. replic.* | A260 | pfu/ml | <A260> | A260_SD | <pfu/ml> | pfu/ml_SD | Ratio | Ratio_SD |
|---|---|---|---|---|---|---|---|---|---|
| T4+_1 | 1 | 10.849 | 8.28E+11 | 10.995 | 0.3078 | 8.43E+11 | 2.46E+11 | 7.66E+10 | 2.25E+10 |
| T4+_1 | 2 | 10.788 | 1.10E+12 | | | | | | |
| T4+_1 | 3 | 11.349 | 6.04E+11 | | | | | | |
| T4+_2 | 1 | 14.93 | 1.20E+12 | 14.627 | 0.2701 | 1.16E+12 | 9.23E+10 | 7.96E+10 | 6.48E+09 |
| T4+_2 | 2 | 14.538 | 1.06E+12 | | | | | | |
| T4+_2 | 3 | 14.412 | 1.24E+12 | | | | | | |
| T4+_3 | 1 | 12.903 | 8.82E+11 | 12.866 | 0.2988 | 9.65E+11 | 7.37E+10 | 7.50E+10 | 5.99E+09 |
| T4+_3 | 2 | 12.55 | 9.92E+11 | | | | | | |
| T4+_3 | 3 | 13.144 | 1.02E+12 | | | | | | |
| 5.4am_1 | 1 | 13.454 | 4.78E+11 | 13.283 | 0.2389 | 5.32E+11 | 4.98E+10 | 4.01E+10 | 3.81E+09 |
| 5.4am_1 | 2 | 13.385 | 5.42E+11 | | | | | | |
| 5.4am_1 | 3 | 13.01 | 5.76E+11 | | | | | | |
| 5.4am_2 | 1 | 12.368 | 5.18E+11 | 12.307 | 0.2151 | 5.06E+11 | 4.72E+10 | 4.11E+10 | 3.90E+09 |
| 5.4am_2 | 2 | 12.485 | 4.54E+11 | | | | | | |
| 5.4am_2 | 3 | 12.068 | 5.46E+11 | | | | | | |
| 5.4am_3 | 1 | 10.45 | 4.34E+11 | 10.471 | 0.0453 | 4.39E+11 | 3.04E+10 | 4.20E+10 | 2.90E+09 |
| 5.4am_3 | 2 | 10.523 | 4.12E+11 | | | | | | |
| 5.4am_3 | 3 | 10.44 | 4.72E+11 | | | | | | |

* A260 and pfu technical replicate measurment are not paired

| Ratio to T4+ (Sample 1) | Scaled SD (to T4+_1) | Sample # |
|---|---|---|
| 1.00E+00 | 2.94E-01 | T4+_1 |
| 1.04E+00 | 8.45E-02 | T4+_2 |
| 9.79E-01 | 7.81E-02 | T4+_3 |
| 5.23E-01 | 4.98E-02 | 5.4am_1 |
| 5.36E-01 | 5.09E-02 | 5.4am_2 |
| 5.47E-01 | 3.79E-02 | 5.4am_3 |

### Figure 5-source data_panel D

**Efficiency of plating (EOP)**

| | | T4+ | | | *5.4am* | | |
|---|---|---|---|---|---|---|---|
| | biol. repli. | used dilution* | pfu count | EOP | used dilution* | pfu count | EOP |
| B178 | 1 | 1.00E-08 | 273 | 0.90 | 1.00E-08 | 335 | 0.95 |
| | 2 | 1.00E-08 | 328 | 1.08 | 1.00E-08 | 361 | 1.03 |
| | 3 | 1.00E-08 | 307 | 1.01 | 1.00E-08 | 360 | 1.02 |
| *hldD ::Tn10* | 1 | 1.00E-08 | 119 | 0.39 | 1.00E-03 | 6 | 1.70E-07 |
| | 2 | 1.00E-08 | 114 | 0.38 | 1.00E-03 | 5 | 1.42E-07 |
| | 3 | 1.00E-08 | 120 | 0.40 | 1.00E-03 | 4 | 1.14E-07 |

* of phage stock, volume was 100µl

### Figure 5-source data_panel E

**Adsorption**

| | biol. repli. | T4+ dilution $10^x$ | T4+ vol (µl) | T4+ pfu count | T4+ pfu/ml* | T4+ adsoprtion (%) | *5.4am* dilution $10^x$ | *5.4am* vol (µl) | *5.4am* pfu count | *5.4am* pfu/ml* | *5.4am* adsoprtion (%) |
|---|---|---|---|---|---|---|---|---|---|---|---|
| LB (ctr) | 1 | -3 | 180 | 113 | 627777.8 | -- | -3 | 180 | 123 | 683333.3 | -- |
| | 2 | -3 | 180 | 117 | 650000.0 | -- | -3 | 180 | 127 | 705555.6 | -- |
| | 3 | -3 | 180 | 86 | 477777.8 | -- | -3 | 180 | 130 | 722222.2 | -- |
| B178 | 1 | -1 | 100 | 27 | 2700.0 | 99.539 | -1 | 100 | 29 | 2900.0 | 99.588 |
| | 2 | -1 | 100 | 30 | 3000.0 | 99.487 | -1 | 100 | 25 | 2500.0 | 99.645 |
| | 3 | -1 | 100 | 22 | 2200.0 | 99.624 | -1 | 100 | 29 | 2900.0 | 99.588 |
| *hldD* ::*Tn10* | 1 | -3 | 180 | 86 | 477777.8 | 18.354 | -3 | 180 | 94 | 522222.2 | 25.789 |
| | 2 | -3 | 180 | 71 | 394444.4 | 32.595 | -3 | 180 | 123 | 683333.3 | 2.895 |
| | 3 | -3 | 180 | 78 | 433333.3 | 25.949 | -3 | 180 | 101 | 561111.1 | 20.263 |

* remaining in the supernatant after centrifugation

### Figure 5-source data_panel F

**phage yield per infected bacteria (burst size)**

| **T4+** | | 5 min -CHCl3 | | | | 5 min + CHCl3 | | | | 30 min -CHCl3 | | | | |
|---|---|---|---|---|---|---|---|---|---|---|---|---|---|---|
| | biol. repli. | dilution $10^x$ | vol (µl) | pfu count | pfu/ml | dilution $10^x$ | vol (µl) | pfu count | pfu/ml | dilution $10^x$ | vol (µl) | pfu count | pfu/ml | **burst size** |
| B178 | 1 | -1 | 50 | 151 | 3.02E+04 | -1 | 400 | 4 | 1.00E+02 | -3 | 25 | 299 | 1.20E+07 | 3.97E+02 |
| | 2 | -1 | 50 | 119 | 2.38E+04 | -1 | 400 | 4 | 1.00E+02 | -3 | 25 | 279 | 1.12E+07 | 4.71E+02 |
| | 3 | -1 | 50 | 132 | 2.64E+04 | -1 | 400 | 5 | 1.25E+02 | -3 | 25 | 284 | 1.14E+07 | 4.32E+02 |
| *hldD* ::*Tn10* | 1 | -1 | 200 | 92 | 4.60E+03 | -1 | 200 | 30 | 1.50E+03 | -3 | 100 | 259 | 2.59E+06 | 8.35E+02 |
| | 2 | -1 | 200 | 112 | 5.60E+03 | -1 | 200 | 32 | 1.60E+03 | -3 | 100 | 162 | 1.62E+06 | 4.05E+02 |
| | 3 | -1 | 200 | 88 | 4.40E+03 | -1 | 200 | 35 | 1.75E+03 | -3 | 100 | 158 | 1.58E+06 | 5.96E+02 |

| ***5.4am*** | | 5 min -CHCl3 | | | | 5 min + CHCl3 | | | | 30 min -CHCl3 | | | | |
|---|---|---|---|---|---|---|---|---|---|---|---|---|---|---|
| | biol. repli. | dilution $10^x$ | vol (µl) | pfu count | pfu/ml | dilution $10^x$ | vol (µl) | pfu count | pfu/ml | dilution $10^x$ | vol (µl) | pfu count | pfu/ml | **burst size** |
| B178 | 1 | -1 | 50 | 140 | 2.80E+04 | -1 | 400 | 2 | 5.00E+01 | -3 | 25 | 160 | 6.40E+06 | 2.29E+02 |
| | 2 | -1 | 50 | 135 | 2.70E+04 | -1 | 400 | 8 | 2.00E+02 | -3 | 25 | 194 | 7.76E+06 | 2.90E+02 |
| | 3 | -1 | 50 | 141 | 2.82E+04 | -1 | 400 | 8 | 2.00E+02 | -3 | 25 | 188 | 7.52E+06 | 2.69E+02 |
| *hldD* ::*Tn10* | 1 | -1 | 200 | 125 | 6.25E+03 | -1 | 200 | 29 | 1.45E+03 | -3 | 100 | 102 | 1.02E+06 | 2.12E+02 |
| | 2 | -1 | 200 | 88 | 4.40E+03 | -1 | 200 | 27 | 1.35E+03 | -3 | 100 | 88 | 8.80E+05 | 2.88E+02 |
| | 3 | -1 | 200 | 81 | 4.05E+03 | -1 | 200 | 30 | 1.50E+03 | -3 | 100 | 105 | 1.05E+06 | 4.11E+02 |

### Figure 5-source data_panel G

**Efficiency of center of infection (ECOI)**

| | | | T4+ | | | | | *5.4am* | | | | |
|---|---|---|---|---|---|---|---|---|---|---|---|---|
| | exp. | biol. repli. | dilution $10^x$ | vol (µl) | pfu count | pfu/ml | ECOI (%) | dilution $10^x$ | vol (µl) | pfu count | pfu/ml | ECOI (%) |
| B178 | 16.03.2015 | 1 | -4 | 25 | 808 | 3.23E+08 | 100.00 | -4 | 25 | 960 | 3.84E+08 | 100.00 |
| | 24.03.2015 | 2 | -4 | 25 | 634 | 2.54E+08 | 98.37 | -4 | 25 | 746 | 2.98E+08 | 95.09 |
| | 24.03.2015 | 3 | -4 | 25 | 655 | 2.62E+08 | 101.63 | -4 | 25 | 823 | 3.29E+08 | 104.91 |
| *hldD* ::*Tn10* | 16.03.2015 | 1 | -4 | 100 | 520 | 5.20E+07 | 16.09 | -3 | 50 | 641 | 1.28E+07 | 3.34 |
| | 24.03.2015 | 2 | -4 | 100 | 314 | 3.14E+07 | 12.18 | -3 | 100 | 578 | 5.78E+06 | 1.84 |
| | 24.03.2015 | 3 | -4 | 100 | 241 | 2.41E+07 | 9.35 | -3 | 100 | 678 | 6.78E+06 | 2.16 |

### Figure 6-source data_panel E

**Adsoprtion and growth of T4+ and T4 *5.4am***

| | T4+ | | | | | | 5.4*am* | | | | | |
|---|---|---|---|---|---|---|---|---|---|---|---|---|
| min | bacteria | dilution 10$^{x}$ | vol (µl) | pfu count | pfu/ml | rel. pfu/ml* | bacteria | dilution 10$^{x}$ | vol (µl) | pfu count | pfu/ml | rel. pfu/ml* |
| 12 | B178 (WT) | -3 | 100 | 1 | 1.00E+04 | 1.15E-02 | B178 (WT) | -3 | 100 | 1 | 1.00E+04 | 1.21E-02 |
| | *hldD* ::Tn*10* | -3 | 200 | 93 | 4.65E+05 | 5.36E-01 | *hldD* ::Tn*10* | -3 | 200 | 91 | 4.55E+05 | 5.53E-01 |
| 60 | B178 (WT) | -4 | 100 | 81 | 8.10E+06 | 9.34E+00 | B178 (WT) | -4 | 100 | 51 | 5.10E+06 | 6.19E+00 |
| | *hldD* ::Tn*10* | -4 | 100 | 160 | 1.60E+07 | 1.85E+01 | *hldD* ::Tn*10* | -3 | 100 | 170 | 1.70E+06 | 2.06E+00 |
| 150 | B178 (WT) | -6 | 200 | 786 | 3.93E+09 | 4.53E+03 | B178 (WT) | -6 | 200 | 268 | 1.34E+09 | 1.63E+03 |
| | *hldD* ::Tn*10* | -5 | 50 | 672 | 1.34E+09 | 1.55E+03 | *hldD* ::Tn*10* | -3 | 50 | 272 | 5.44E+06 | 6.61E+00 |

* relative to phage inoculum

| phage inoculum | pfu/ml |
|---|---|
| T4+ | 8.67E+05 |
| 5.4*am* | 8.23E+05 |

### Figure S2-source data_panel A

**Efficiency of plating (EOP) of T4+ and T4 *5.4am***

| | T4+ | | | | | 5.4*am* | | | | |
|---|---|---|---|---|---|---|---|---|---|---|
| | biol. repli. | vol (µl)* | pfu count | pfu/ml | relative EOP | biol. repli. | vol (µl)* | pfu count | pfu/ml | relative EOP |
| K12 sup+ (B178) | 1 | 100 | 174 | 1.74E+03 | 1.10E+00 | 1 | 100 | 155 | 1.55E+03 | 1.33E+00 |
| | 2 | 100 | 181 | 1.81E+03 | 1.14E+00 | 2 | 100 | 141 | 1.41E+03 | 1.21E+00 |
| | 3 | 100 | 191 | 1.91E+03 | 1.21E+00 | 3 | 100 | 132 | 1.32E+03 | 1.13E+00 |
| K12 supF (NM538) | 1 | 100 | 163 | 1.63E+03 | 1.03E+00 | 1 | 100 | 120 | 1.20E+03 | 1.03E+00 |
| | 2 | 100 | 172 | 1.72E+03 | 1.09E+00 | 2 | 100 | 125 | 1.25E+03 | 1.07E+00 |
| | 3 | 100 | 140 | 1.40E+03 | 8.84E-01 | 3 | 100 | 104 | 1.04E+03 | 8.94E-01 |
| K12 supD (CR63) | 1 | 100 | 200 | 2.00E+03 | 1.26E+00 | 1 | 100 | 142 | 1.42E+03 | 1.22E+00 |
| | 2 | 100 | 181 | 1.81E+03 | 1.14E+00 | 2 | 100 | 127 | 1.27E+03 | 1.09E+00 |
| | 3 | 100 | 155 | 1.55E+03 | 9.79E-01 | 3 | 100 | 113 | 1.13E+03 | 9.71E-01 |
| B sup+ (BE) | 1 | 100 | 230 | 2.30E+03 | 1.45E+00 | 1 | 100 | 167 | 1.67E+03 | 1.44E+00 |
| | 2 | 100 | 214 | 2.14E+03 | 1.35E+00 | 2 | 100 | 141 | 1.41E+03 | 1.21E+00 |
| | 3 | 100 | 201 | 2.01E+03 | 1.27E+00 | 3 | 100 | 167 | 1.67E+03 | 1.44E+00 |

* For each phage strain ( T4+ and T4 5.4am), a single ~1.5*103 pfu/ml dilution was used to assess pfu on the indicated bacterial strains

### Figure S2-source data_panel B

**Stability of T4+ and T4 *5.4am***

| | T4+ | | | | | | 5.4*am* | | | | | |
|---|---|---|---|---|---|---|---|---|---|---|---|---|
| day | biol. repli. | dilution $10^x$ | vol (µl) | pfu count | pfu/ml | rel. pfu/ml | biol. repli. | dilution $10^x$ | vol (µl) | pfu count | pfu/ml | rel. pfu/ml |
| 0 | 1 | -2 | 100 | 212 | 2.12E+05 | 9.44E-01 | 1 | -2 | 100 | 192 | 1.92E+05 | 9.16E-01 |
| | 2 | -2 | 100 | 237 | 2.37E+05 | 1.06E+00 | 2 | -2 | 100 | 227 | 2.27E+05 | 1.08E+00 |
| 1 | 1 | -2 | 100 | 117 | 1.17E+05 | 5.21E-01 | 1 | -2 | 100 | 107 | 1.07E+05 | 5.11E-01 |
| | 2 | -2 | 100 | 130 | 1.30E+05 | 5.79E-01 | 2 | -2 | 100 | 131 | 1.31E+05 | 6.25E-01 |
| 3 | 1 | -1 | 50 | 210 | 4.20E+04 | 1.87E-01 | 1 | -1 | 50 | 218 | 4.36E+04 | 2.08E-01 |
| | 2 | -1 | 50 | 227 | 4.54E+04 | 2.02E-01 | 2 | -1 | 50 | 210 | 4.20E+04 | 2.00E-01 |
| 6 | 1 | -1 | 200 | 180 | 9.00E+03 | 4.01E-02 | 1 | -1 | 200 | 223 | 1.12E+04 | 5.32E-02 |
| | 2 | -1 | 200 | 188 | 9.40E+03 | 4.19E-02 | 2 | -1 | 200 | 212 | 1.06E+04 | 5.06E-02 |

### Figure S2-source data_panel C

##### Adsorption rate of T4+ and T4 *5.4am*

| | T4+ | | | | | 5.4*am* | | | | |
|---|---|---|---|---|---|---|---|---|---|---|
| min | dilution $10^x$ | vol (µl) | pfu count | pfu/ml | rel. pfu/ml | dilution $10^x$ | vol (µl) | pfu count | pfu/ml | rel. pfu/ml |
| 0 | -3 | 50 | 179 | 3.58E+06 | 1.00E+00 | -3 | 50 | 167 | 3.34E+06 | 1.00E+00 |
| 1 | -3 | 50 | 110 | 2.20E+06 | 6.15E-01 | -3 | 50 | 112 | 2.24E+06 | 6.71E-01 |
| 2 | -3 | 50 | 76 | 1.52E+06 | 4.25E-01 | -3 | 50 | 67 | 1.34E+06 | 4.01E-01 |
| 3 | -3 | 50 | 41 | 8.20E+05 | 2.29E-01 | -3 | 50 | 46 | 9.20E+05 | 2.75E-01 |
| 4 | -3 | 100 | 74 | 1.48E+06 | 4.13E-01 | -3 | 100 | 62 | 1.24E+06 | 3.71E-01 |
| 6 | -3 | 100 | 34 | 6.80E+05 | 1.90E-01 | -3 | 100 | 33 | 6.60E+05 | 1.98E-01 |
| 8 | -2 | 50 | 93 | 1.86E+05 | 5.20E-02 | -2 | 50 | 87 | 1.74E+05 | 5.21E-02 |
| 10 | -2 | 50 | 47 | 9.40E+04 | 2.63E-02 | -2 | 50 | 59 | 1.18E+05 | 3.53E-02 |
